# Calponin-Homology Domain mediated bending of membrane associated actin filaments

**DOI:** 10.1101/2020.07.10.197616

**Authors:** Saravanan Palani, Andrejus Suchenko, Sayantika Ghosh, Esther Ivorra-Molla, Scott Clarke, Mohan K. Balasubramanian, Darius V. Köster

**Affiliations:** Centre for Mechanochemical Cell Biology and Warwick Medical School, Division of Biomedical Sciences, CV4 7AL Coventry, UK; Department of Biochemistry, Division of Biological Sciences, Indian Institute of Science, Bangalore-560012, India

## Abstract

Actin filaments are central to numerous biological processes in all domains of life. Driven by the interplay with molecular motors, actin binding and actin modulating proteins, the actin cytoskeleton exhibits a variety of geometries. This includes structures with a curved geometry such as axon-stabilizing actin rings, actin cages around mitochondria and the cytokinetic actomyosin ring, which are generally assumed to be formed by short linear filaments held together by actin cross-linkers. However, whether individual actin filaments in these structures could be curved and how they may assume a curved geometry remains unknown. Here, we show that “curly”, a region from the IQGAP family of proteins from three different organisms, comprising the actin-binding calponin-homology domain and a C-terminal unstructured domain, stabilizes individual actin filaments in a curved geometry when anchored to lipid membranes. Whereas F-actin is semi-flexible with a persistence length of ~10 μm, binding of mobile curly within lipid membranes generates actin filament arcs and full rings of high curvature with radii below 1 μm. Higher rates of fully formed actin rings are observed in the presence of the actin-binding coiled-coil protein tropomyosin and also when actin is directly polymerized on lipid membranes decorated with curly. Strikingly, curly induced actin filament rings contract upon the addition of muscle myosin II filaments and expression of curly in mammalian cells leads to highly curved actin structures in the cytoskeleton. Taken together, our work identifies a new mechanism to generate highly curved actin filaments, which opens a new range of possibilities to control actin filament geometries, that can be used, for example, in designing synthetic cytoskeletal structures.

---

The IQGAP family of proteins plays a key role in actin cytoskeleton regulation including the assembly and function of the contractile actomyosin ring in budding and fission yeasts (Briggs & Sacks, 2003; Eng et al., 1998; Epp & Chant, 1997; Lippincott & Li, 1998; Tebbs & Pollard, 2013). To study the mechanism and role of actin binding by the fission yeast IQGAP (encoded by the *rng2* gene), we utilized a strategy to investigate its function when immobilized on supported lipid bilayers. We chose this approach, since during cytokinesis Rng2, which binds several actomyosin ring proteins, is tethered to the plasma membrane via Mid1, ensuring the formation and anchoring of the cytokinetic ring (Laplante et al., 2016; Laporte et al., 2011; Padmanabhan et al., 2011). We linked hexa-histidine tagged *rng2* protein fragments to supported lipid bilayers containing nickel-chelating lipids (DOGS-NTA(Ni^2+^)) and observed the binding of fluorescently labelled actin filaments using live total internal reflection fluorescence (TIRF) microscopy as described earlier (Köster et al., 2016) (Figure 1A). The actin-binding calponin homology domain (CHD) is located at the N-terminus of Rng2 (AA 41-147), and the construct His_6_-Rng2(1-189) (subsequently referred to as curly) containing the CHD and additional 42 amino acids was found to bind actin filaments with K_d_ = 0.9 ± 0.3 μM (Figure 1B, Figure 1-figure supplement 1 A-C), as reported in (Hayakawa et al., 2020). Remarkably, in presence of His_6_-Curly, most actin filaments settling and binding onto the SLB formed tightly bent arcs and full rings with curvatures of C_curly_ = 1.7 ± 0.5 μm^−1^ (Figure 1 B, C; Figure 1 – figure supplement 1 D-G; Video 1). To our knowledge, this is an unprecedented phenomenon specific to curly. Binding of other membrane tethering actin binding proteins in the same geometry, such as the CHD of α-actinin or the actin binding domain of Ezrin, did not appreciably bend actin filaments (Figure 1-figure supplement 2 A-D). Membrane anchored fimbrin has also been shown not to bend actin (Murrell & Gardel, 2012).

**Figure 1.**
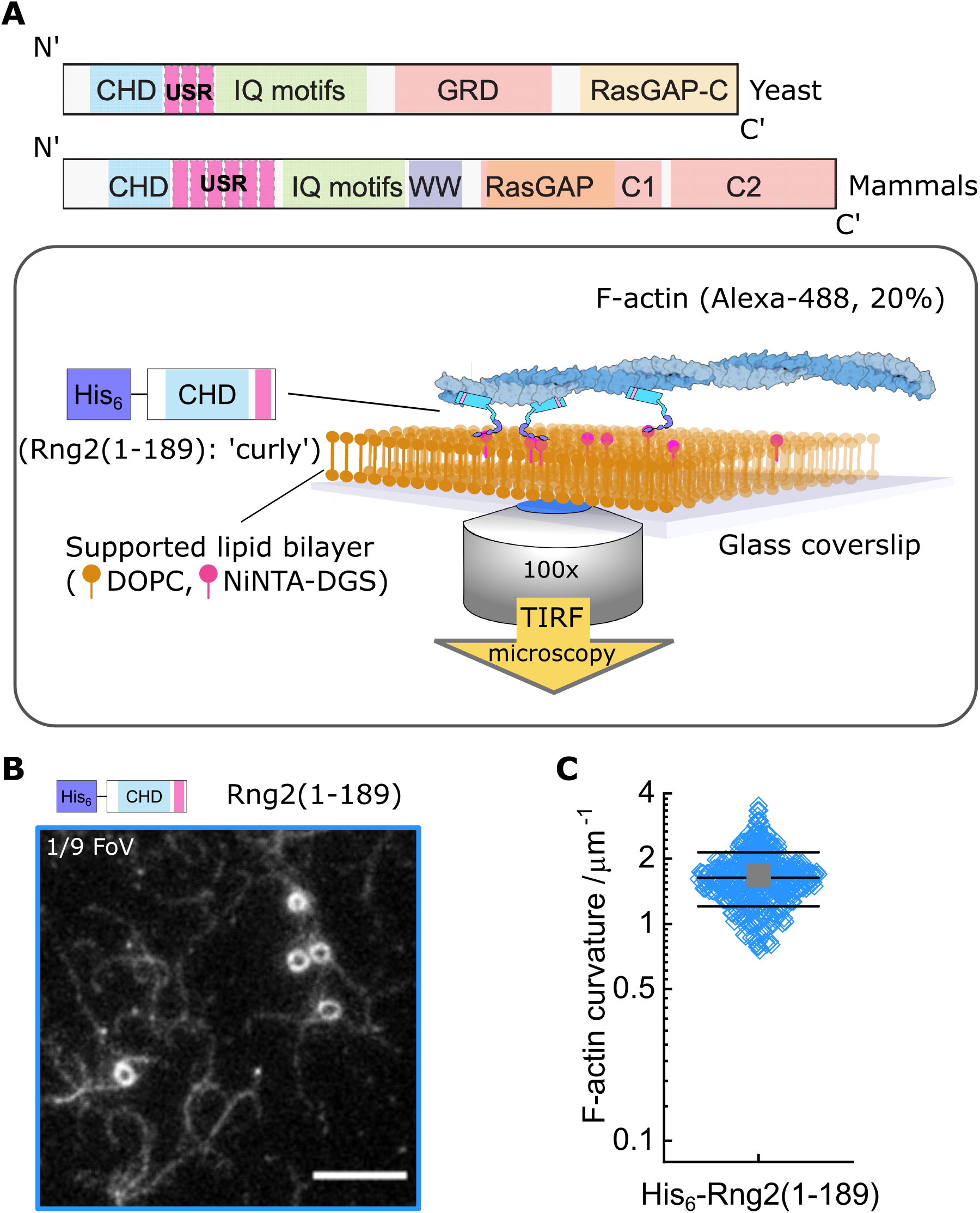
Formation of actin filament rings by membrane tethered curly (Rng2(1-189)) (A) Schematic representation of (top) the IQGAP proteins Rng2 (yeast, S. pombe) and IQGAP1 (mammals, H. Sapiens) and (bottom) the experimental setup used in this study; CHD - Calponin Homology Domain, USR – Unstructured Region, GRD - GAP Related Domain, RasGAP – Ras GTPase Activating Protein, WW – tryptophan containing protein domain. (B) TIRF microscopy image of actin filaments (Alexa488, Cactin = 100 nM) bound to SLB tethered His_6_-curly (C_curly_ = 10 nM); shown is 1/9 field of view (FoV), scale bar 5 μm. (C) Curvature measurements of actin filament rings and curved segments; shown are the individual data points and their mean ± s.d.; N = 425 obtained from 5 field of views from each of 4 independent experiments.

To understand the mechanism leading to actin filament bending and ring formation by curly, we tested the role of different fragments of curly, their orientation, and curly anchoring to lipid membranes in actin filament bending. Curly mobility within planar lipid membranes was important for actin bending as glass adsorbed, immobilized His_6_-Curly displayed reduced actin binding and bending (C_glass_ = 0.6 ± 0.3 μm^−1^) (Figure 2 A). Next, we generated fragments of curly to discern the regions important for actin binding and bending. We found that the C-terminal region following the CHD alone, His_6_-Rng2(150-250), did bind actin filaments without inducing bending (C_(150-250)_ = 0.2 ± 0.1 μm^−1^) (Figure 2 B). Interestingly, a 7AA deletion, Rng2(1-189)Δ154-160, led to a reduced degree of actin binding and bending (CΔ(154-160) = 0.4 ± 0.1 μm^−1^) (Figure 2 C). Similarly, the fragments Rng2(41-189) and Rng2(1-147) displayed weaker actin binding and bending compared to curly (C_(41-189)_ = 0.6 ± 0.2 μm^−1^; C_(1-147)_ = 0.7 ± 0.2 μm^−1^) (Figure 2 D, E). Changing the location of the hexa-histidine tag to the C-terminus, Rng2(1-189)-His_6_, did not affect actin filament bending (C_curly-his_ = 1.5 ± 0.5 μm^−1^) (Figure 2 F, Figure 2-figure supplement 1 C). Taken together (Figure 2 G; Figure 2-figure supplement 1 A, B), this suggests that curly contains two actin binding sites, one located within the CHD followed by a second within Rng2(148-189). Both actin binding sites are necessary for actin bending as neither Rng2(1-147) nor Rng2(150-250) caused strong bending. The second actin binding site likely includes the 7AA Rng2(154-160) as this unstructured region maps directly to AA 240-246 of the dystrophin CH2 domain (Wang et al., 2004). The first 40 AA of Rng2 are likely to be important for the protein folding and stability, because of which the Rng2(41-189) construct showed poor actin binding and did not lead to actin bending.

**Figure 2.**
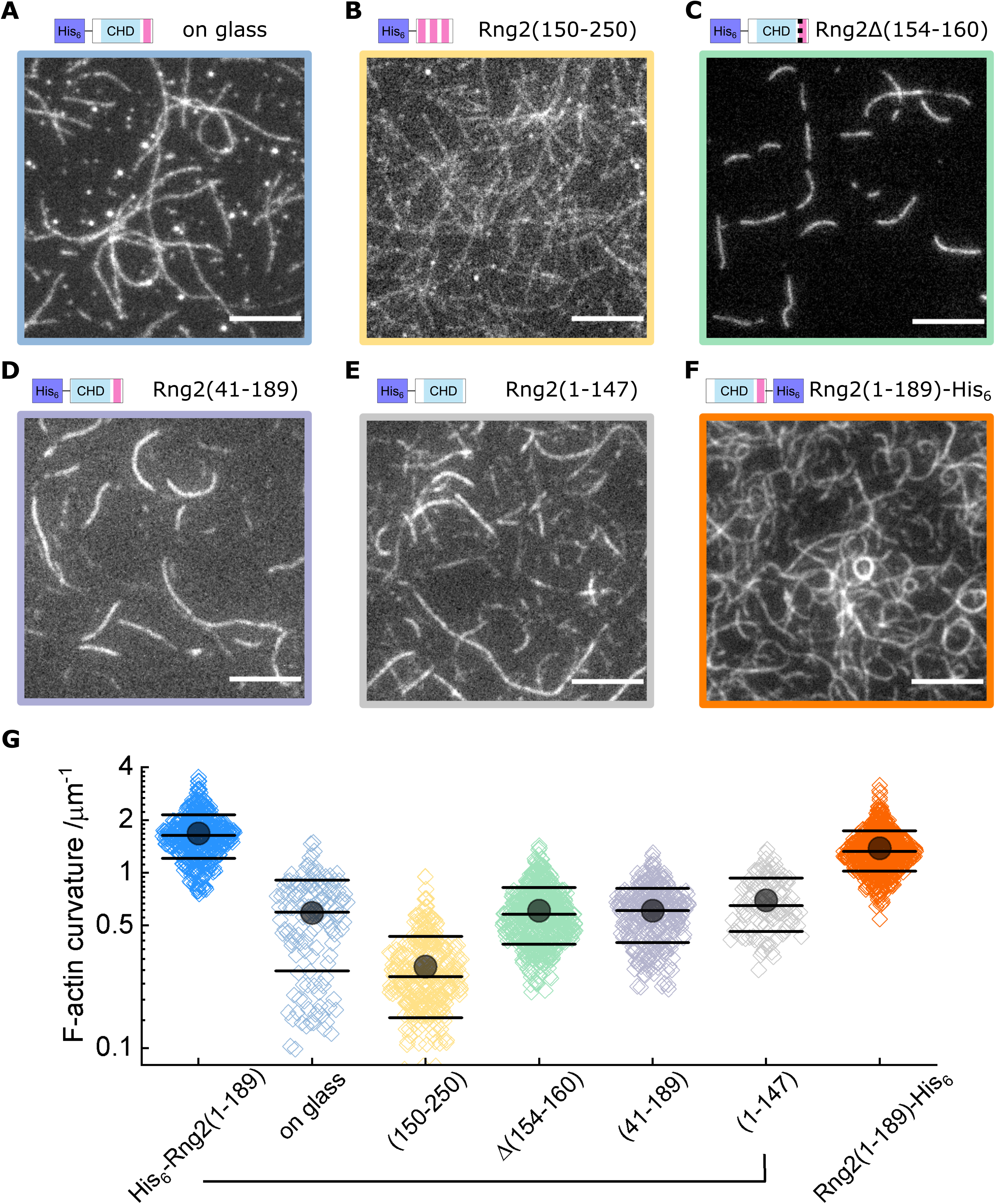
Characterization of actin bending by fragments of curly. TIRF microscopy images of actin filaments (Alexa488, C_actin_ = 100 nM) bound to (A) glass adsorbed His_6_-curly (Ccurly = 10 nM); N = 138 from 9 field of views from each of 3 independent experiments; (B) SLB bound His_6_-Rng2(150-250)(C_(150-250)_ = 10 nM); N = 327 from 20 field of views from each of 2 independent experiments; (C) SLB bound His_6_-Rng2(1-189)Δ154-160(C_Δ(154-160)_ = 10 nM); N = 593 from 15 field of views from each of 3 independent experiments; (D) SLB bound His_6_-Rng2(41-189)(C(41-189) = 10 nM); N = 323 from 20 field of views from each of 3 independent experiments; (E) SLB bound His_6_-Rng2(1-147)(C_(1-147)_ = 10 nM); N = 118 from 12 field of views from each of 3 experiments; (F) SLB bound Rng2(1-189)-His_6_(CCurly-His = 10 nM); N = 658 from 13 field of views from each of 4 experiments; each image shows 1/9 field of view (FoV); scale bars: 5 μm. (G) Curvature measurements of actin filament rings and curved segments; diamonds represent individual measurements, lines the median ± standard deviation and the circle the mean value.

Next, we studied whether actin bending by curly depended on the orientation of actin filaments by following the landing of already polymerized actin filaments decorated with labelled capping protein as a plus end marker (Bieling et al., 2016). We found that the bending was oriented anti-clockwise with respect to the plus end in all instances, wherein the plus end was clearly labelled to identify the orientation of filament bending (Figure 3A, B; Figure 3-figure supplement 1 A, B). This was observed using both, the N-terminal and C-terminal hexa-histidine tagged curly, indicating that the internal sequence of the two actin binding sites within curly sets the chirality of actin bending and not the position of the membrane linker (Figure 3A, B; Figure 3-figure supplement 1 A, B; Video 3, 4). Actin filaments appeared to bend concomitant with their landing on the supported lipid bilayer, which indicates that the bending did not require the full actin filament to be tethered to the SLB and underlined the earlier observation that the bending occurred locally.

**Figure 3.**
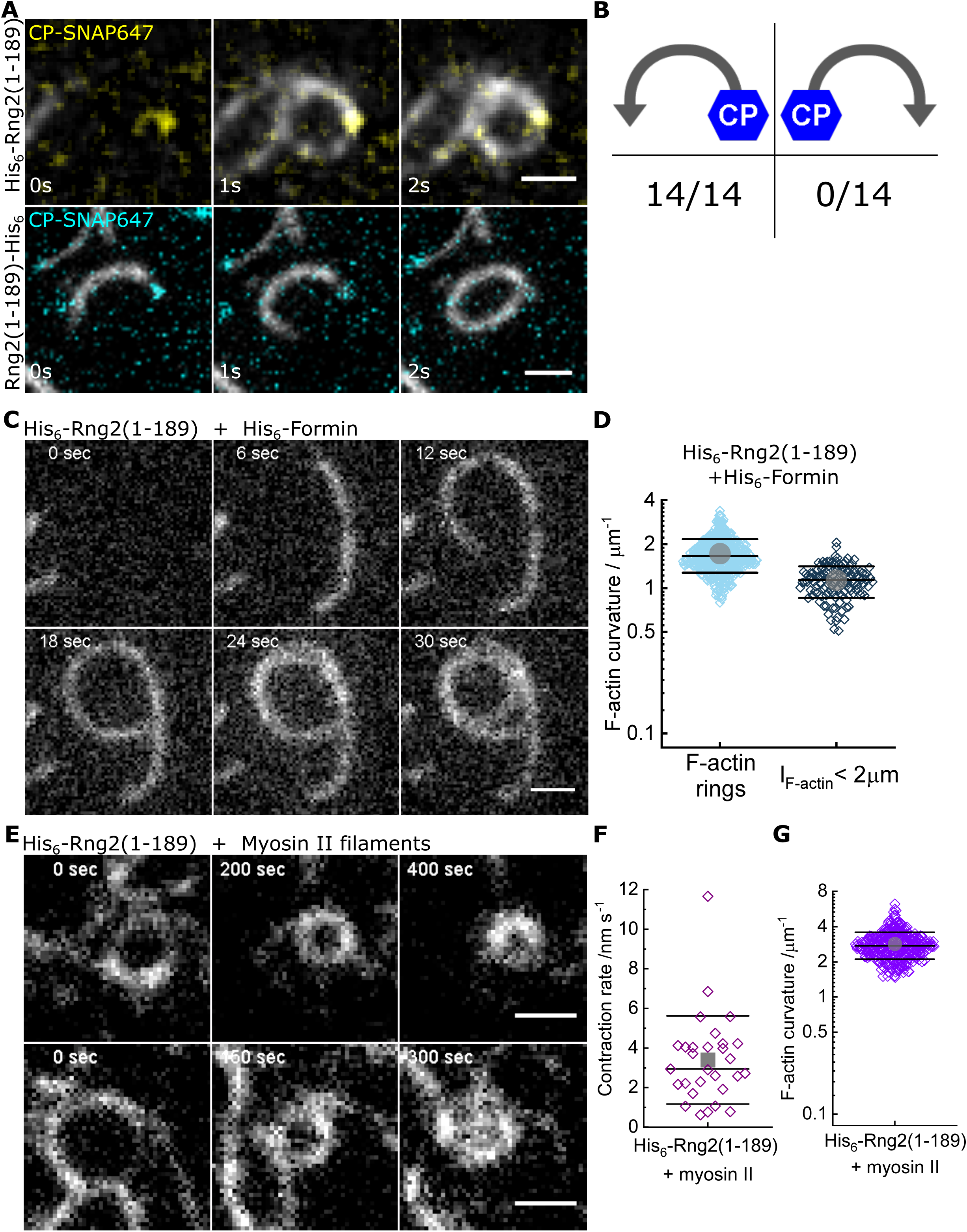
Curly recognizes actin filament orientation and enables actin ring contraction by myosin II. (A) TIRF microscopy images of actin filaments (Alexa488) (C_actin_ = 100 nM) with the plus end marked with SNAP647-tagged capping protein (CCP = 2 nM) binding to His_6_-curly (top) and curly-His_6_ (bottom) (C_curly_ = 10 nM); scale bar: 1 μm. (B) Count of actin filament bending orientations with respect to the capping protein where individual actin filaments could be identified. (C) TIRF microscopy images of a polymerizing actin filament (Alexa488) driven by membrane tethered His_6_-formin in the presence of His_6_-curly; scale bar: 1 μm. (D) Curvature measurements of actin filament rings (light blue) and curved short actin filaments (< 2 μm; grey-blue); shown are the individual data points and their mean ± s.d.; Nrings = 477, Nshort = 125 from 9 field of views of 3 independent experiments. (E) TIRF microscopy images of actin filament (Alexa488) ring contraction after addition of rabbit muscle myosin II filaments on His_6_-curly containing SLBs; scale bar: 1 μm. (F) Average contraction rates of actin filament rings after addition of rabbit muscle myosin II filaments; shown are the individual data points and their mean ± s.d.; N = 18 from 2 individual experiments. (G) Curvature measurements of actin filament rings and curved segments 20 min after addition of rabbit muscle myosin II filaments; shown are the individual data points and their mean ± s.d.; N = 342 from 10 field of views of 2 individual experiments.

To decouple the actin filament bending from the landing of actin filaments, we induced the polymerization of new actin filaments at planar lipid membranes by membrane tethered formin (His_6_-SpCdc12(FH1-FH2)), profilin-actin and ATP in the presence of membrane tethered His_6_-Curly. Strikingly, actin filaments displayed characteristic bending shortly after the onset of polymerization (C_formin, short_ = 1.1 ± 0.3 μm^−1^) and often grew into full rings (C_formin rings_ = 1.7 ± 0.4 μm^−1^) (Figure 3 C, D; Figure 3-figure supplement 2 A-E; Video 5). By contrast, polymerization of actin filaments along SLBs decorated with His_10_-SNAP-EzrinABD did not result in the formation of arcs or rings, establishing that actin filament bending was due to curly and not due to formin (Figure 3-figure supplement 2 F, G). These observations show that actin bending occurs continuously due to the binding of membrane tethered curly and did not require the cross-linking of adjacent ends of the same filament as was observed with the actin cross-linker anillin (Kučera et al., 2020). Importantly, the uni-directional bending supports the hypothesis that the binding site of curly with actin filaments defines an orientation with the propagation of an established curved trajectory indicating a cooperative process.

Actin filaments forming the cytokinetic ring in *S. pombe* are tightly associated with tropomyosin (Cdc8). In contrast, the actin cross-linker fimbrin is present outside the cytokinetic ring region in Arp2/3 generated actin patches and prevents tropomyosin binding to these patches (Skau & Kovar, 2010). To determine whether the actin bending effect of curly is conserved in tropomyosin-wrapped actin filaments, we incubated actin filaments with tropomyosin before adding them to His_6_-Curly containing SLBs. Strikingly, addition of tropomyosin to actin filaments increased the frequency of actin ring formation without affecting actin filament curvature, while actin filaments incubated with the actin cross-linker fimbrin displayed reduced bending and ring formation (Figure 3-figure supplement 3; Video 6). Thus, the tropomyosin Cdc8 and curly cooperate to enhance actin filament bending and ring formation.

Interestingly, we could observe that long actin filaments coated with tropomyosin would trace consecutive rings around the same center while landing on curly decorated lipid membranes. Subtraction of the image after completion of the first round of actin filament landing into a ring from the image after the second round revealed that the second ring occupied the interior space of the first ring. In line with that, a comparison of the intensity profiles perpendicular to the actin filament of the first and second round of ring formation revealed a widening of the profile towards the ring’s interior (Figure 3-figure supplement 4 A, B). A similar effect could be observed in examples of actin filaments polymerized by membrane tethered formin in the presence of membrane tethered curly (Figure 3-figure supplement 4 C, D). This would suggest that curly can arrange long actin filaments into an inward-oriented spiral.

To test whether the curly-induced actin rings can contract, we added rabbit skeletal muscle myosin II filaments and ATP to actin filaments bound to SLB tethered His_6_-curly and followed actin filament dynamics over time. Shortly after myosin II addition, actin filaments (curved and straight ones) were propelled by myosin action leading to increased bending, rotation and finally to ring formation and contraction (Figure 3 E-G, Figure 3-figure supplement 5 A-D, Video 7). Interestingly, most actin rings displayed a counter-clockwise rotation (34/36 cases) and a slow contraction speed of 3 ± 2 nm s^−1^ (Figure 3 F; Video 8). The density of actin rings was strongly increased after myosin II addition (Figure 3-figure supplement 5E), indicating that actin sliding leads to more recruitment of curly. In line with this, actin filament rings displayed increased localization of fluorescently labelled curly after addition of myosin II filaments (Figure 3-figure supplement 5 F-H). Despite reaching very high actin curvatures (up to 6.3 μm^−1^) no breaking of actin filaments during the contraction process could be observed suggesting that binding of curly reduces the rigidity of actin filaments.

Since Rng2 belongs to the IQGAP protein family, we tested the N-terminal hexa-histidine tagged fragments of the IQGAP proteins Iqg1(1-330) (*S. cerevisiae*) and IQGAP1(1-678) (*H. sapiens*) and found that the bending of actin filaments was conserved (C_S.C._ = 1.1 ± 0.4 μm^−1^; C_H.S._ = 1.0 ± 0.2 μm^−1^) (Figure 4 A-C; Figure 4-figure supplement 1 A). Comparison of the available crystal structures of *H. sapiens* IQGAP1(28-190) with *S. pombe* Rng2(32-190) indicates high similarity between the two (Figure 4-figure supplement 1 B).

**Figure 4.**
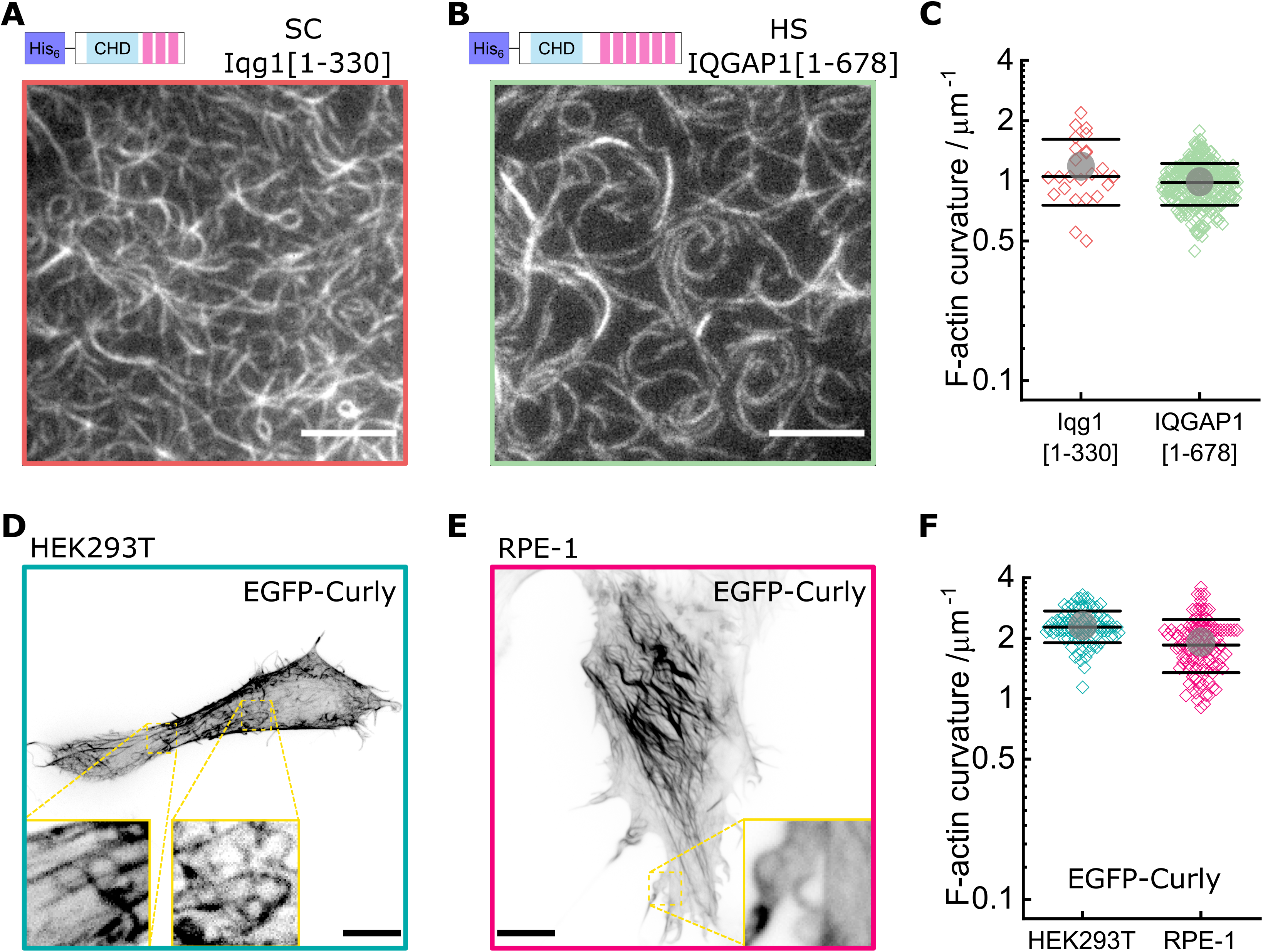
Curly effect is conserved among species and can foster actin bending in mammalian cells. (A) TIRF microscopy image of actin filaments (Alexa488) bound to membrane tethered His_6_-Iqg1(1-330) (S. cerevisiae); image shows 1/9 of the field of view, scale bar: 5 μm. (B) TIRF microscopy image of actin filaments (Alexa488) bound to membrane tethered His_6_-IQGAP1(1-678) (H. sapiens); image shows 1/9 of the field of view, scale bar: 5 μm. (C) Curvature measurements of actin filament rings and curved segments; shown are the individual data points and their mean ± s.d.; Iqg1(1-330) (orange): N = 110 from 3 field of views of each of 3 individual experiments; IQGAP1(1-678) (green): N = 407 from 20 field of views of 3 individual experiments. (D) Confocal microscopy image (maximum intensity projection of the basal cell section) of a HEK293T cell transfected with EGFP-Rng2(1-189), inlet shows zoom of dashed box; scale bar: 5 μm. (E) Confocal microscopy image (maximum intensity projection of the basal cell section) of a RPE-1 cell transfected with EGFP-Rng2(1-189), inlet shows zoom of dashed box; scale bar: 5 μm. (F) Curvature measurements of actin filament rings and curved segments found in EGFP-Rng2(1-189) expressing cells; shown are the individual data points and their mean ± s.d.; HEK293T (teal): N 91 from 14 cells of 2 independent experiments; REP-1 (fuchsia): N = 113 from 11 cells of 2 independent experiments.

Finally, to test the effect of curly on the actin cortex in cells, we expressed GFP tagged curly in the mammalian cell lines HEK293T and RPE-1, which resulted in striking changes of the actin cortex architecture with the prominent occurrence of curved and ring-shaped actin filaments and bundles with curvatures of C_HEK293T_ = 2.3 ± 0.4 μm^−1^ and C_RPE-1_ = 1.9 ± 0.6 μm^−1^ (Figure 4D-F, Figure 4-figure supplement 2 A-D, Figure 4-figure supplement 3). Co-expression with LifeAct-mCherry confirmed EGFP-Rng2(1-189) bound to actin filaments in cells (Figure 4-figure supplement 2 A, B). To our surprise, addition of a CaaX domain to tether curly to cellular membranes did not enhance the actin bending effect but located curly mainly to structures resembling the endoplasmic reticulum (Figure 4-figure supplement 4 A). The EGFP-Rng2(144-189) and EGFP-Rng2(144-189)-CaaX constructs showed only cytoplasmic localization (Figure 4-figure supplement 4 B, C). To test whether the actin bending effect is conserved in the mammalian full-length protein, we imaged HEK293T cells expressing full length EGFP-IQGAP1 and LifeAct-mCherry using lattice light sheet microscopy and indeed observed curved and ring-shaped actin structures inside the cell as well at its surface (Figure 4-figure supplement 5). These experiments established that curly could instructively reorganize actin filaments into curved structures and rings and that this capacity is conserved in full-length IQGAP1.

## Discussion

Our results show that the N-terminal CHD of IQGAP proteins induces actin filament bending when tethered to lipid membranes, which constitutes a new type of actin binding protein and could be an important link between actin and membrane geometries. CHDs of other proteins, such as α-actinin (K_d_ = 4.7 μM (Wachsstock et al., 1993)) or utrophin (Kd = 1.5 μM (Singh et al., 2017)) do not show this behavior. Recently, Uyeda and colleagues reported that curly (Rng2(1-189)) in solution could induce kinks at random locations of the actin filament (Hayakawa et al., 2020). When constrained to a lipid membrane, asymmetric binding of curly to actin filaments could lead to a succession of kinks towards the same direction leading to continuous actin filaments bending into rings in the plane of the lipid membrane. With an estimated His_6_-Curly surface density on SLBs of 5000 μm^−2^ (Köster et al., 2016; Nye & Groves, 2008) the approximated curly to actin ratio would be 1:7 or higher. The mobility of curly on the SLB allowing accumulation under actin filaments is essential for actin filament bending as glass-immobilized curly failed to generate rings. Based on the data of the Utrophin-CHD actin binding sites (Kumari et al., 2020), the newly actin binding region within Rng2(148-189) identified here, suggest binding to the actin D-loop and could reduce the effective actin persistence length below its typical value of 10 μm (De La Cruz & Gardel, 2015; Mccullough et al., 2008). The increased flexibility of actin filaments is highlighted by the fact that addition of rabbit muscle myosin II filaments resulted in actin ring constriction without any evidence for filament rupture up to curvatures of 6.3 μm^−1^, which is much higher than actin alone (Taylor et al., 2000). This mechanism of actin ring formation stands out as it bends individual actin filaments in contrast to other reported systems that generate actin rings made of bundles of actin filaments (Litschel et al., 2020; Mavrakis et al., 2014; Mishra et al., 2013; Miyazaki et al., 2015; Way et al., 1995).

It was not evident that the addition of myosin II filaments would lead to actin ring constriction without the addition of any cross-linkers or other factors. When considering that curly arranges actin filaments into an inward spiral, a possible explanation for actin ring constriction would be that the myosin II filament acts both as a cross-linker and motor protein: one end of the myosin II filament sits at the actin filament plus end while other myosin head domains of the same myosin II filament pull along the same actin filament to travel towards the plus end leading to constriction. This would result in sub-optimal myosin head orientations towards the actin filament, which could explain the observed slow constriction rates that were orders of magnitude slower than the reported values for actin propulsion by myosin II in motility assays (Toyoshima et al., 1990). Future studies will hopefully provide more insights into this peculiar minimal ring constriction mechanism.

Highly bent actin filament structures are most likely important for many cellular structures such as axons (Vassilopoulos et al., 2019; Xu et al., 2013) and mitochondrial actin cages (Kruppa et al., 2018), but the molecular mechanisms leading to their formation are still poorly understood. Future work could provide insights into whether curly plays a role in actin ring formation in axons and around mitochondria. In addition, our system of membrane-bound curly, actin filaments, and myosin II filaments constitutes a minimalistic system for actin ring formation and constriction and could be used in the future to design synthetic dividing vesicles and further exiting active membrane-cortex systems.

## Materials and Methods

### Cloning and Protein purification

S. pombe Rng2 fragments, Fim1, Cdc12 (FH1-FH2) and S. cerevisiae Iqg1 were amplified from cDNA library and genomic DNA respectively. Amplified fragments were cloned into pET (His_6_) and pGEX (GST) based vectors using Gibson cloning method (NEB builder, E5520S). Plasmids used in this study is listed in Table S1.

All protein expression plasmids were transformed into E.coli BL21-(DE3). Single colony was inoculated in 20□ml of LB media supplemented with appropriate antibiotic (pET-Kanamycin; pGEX-Ampicillin). Precultures were grown for ∼12-16□h at 36□°C shaking at 20□r.p.m. Cells were diluted to OD600 of 0.1□a.u. in 500□ml of LB with antibiotics and protein expression was induced with 0.25□mM isopropyl β-D-1-thiogalactopyranoside (IPTG). Protein was expressed for 3-4□h at 30□°C shaking at 200□r.p.m. unless otherwise noted. After induction cell pellets were collected and spun down at 7,000 r.p.m for 20 minutes after induction at 4□°C. Media was aspirated and pellets were washed once with cold phosphate buffered saline (PBS) with 1mM phenylmethylsulfonyl fluoride (PMSF), and pellets were stored at −80□°C.

Ni-NTA purification of truncated IQGAP proteins (His_6_ tagged Rng2, Iqg1 and IQGAP1): Cell pellets for purification were thawed on ice for 10 minutes. The pellets were resuspended in 10□ml of lysis buffer for sonication (50□mM Napi pH 7.6, 200□mM NaCl, 10mM Imidazole pH 7.5, 0.5□mM EDTA, 1□mM DTT, 1□mg/ ml lysozyme, and complete mini-EDTA-Free protease inhibitor cocktail) and incubated on ice for 20□min, followed by sonication (8 cycles, 15 sec pulse). The lysates were centrifuged at 14000 r.p.m, 30□min, 4□°C and the clarified lysate was transferred to a 15-ml tube. The 400□μl slurry of HisPur™ Ni-NTA agarose resin (cat. no. 88221, Thermo fisher) was washed with wash buffer (5x) (50□mM Napi (pH 7.6), 300□mM NaCl, 30mM Imidazole pH 7, 0.5□mM EDTA and 1□mM DTT) before the lysate was added. The clarified lysate was added to the washed Ni-NTA resin and incubated for 2h at 4□°C. After incubation with NiNTA resin, beads were washed with wash buffer 6-8 times in poly-prep chromatography columns (cat. no. 7311550, BIO-RAD laboratories Inc). Protein was eluted using Ni-NTA elution buffer (50□mM NaPi pH 7.6, 300□mM NaCl, 0.5□mM EDTA, 1□mM DTT and 500□mM imidazole) and 300□μl elutions were collected in a clean Eppendorf tubes. Each fraction was assessed by SDS– polyacrylamide gel electrophoresis (SDS–PAGE). The eluates (E1-E3) were pooled, concentrated and buffer exchanged into the protein storage buffer (50□mM Tris-HCl pH 7.4, 150□mM NaCl, 1□mM DTT and 10% glycerol) using a PD MiniTrap G-25 sephadex columns (GE Healthcare) and the purified proteins were flash frozen in liquid N_2_ and stored at −80□°C. The protein concentration was estimated by UV280 and by comparing known quantities of BSA standards on an SDS–PAGE gel.

GST tagged protein (GST-Fim1) purification: Cell pellets for purification were thawed on ice for 10 minutes. The pellets were resuspended in 10□ml of lysis buffer for sonication (PBS, 0.5□mM EDTA, 1□mM DTT, 1□mg/ ml lysozyme, and complete mini-EDTA-Free protease inhibitor cocktail tablets) and incubated on ice for 20□min, followed by sonication (10 cycles, 15 sec pulse). After sonication cell lysate was incubated with 1% Triton-X-100 for 20 minutes on ice. The lysates were centrifuged at 22000xg, 30□min, 4□°C and the clarified lysate was transferred to a 15-ml tube. The 400□μl slurry of glutathione sepharose-4B resin (cat. no. GE17-0756-01, GE) was washed with wash buffer (5x) (PBS, 0.5□mM EDTA and 1□mM DTT) before the lysate was added. The clarified lysate was added to the washed glutathione sepharose resin and incubated for 2-3h at 4□°C. After incubation with sepharose resin, beads were washed with wash buffer 6-8 times in poly-prep chromatography columns. Protein was eluted using GST elution buffer (50□mM Tris-HCl pH8.0 and 10□mM glutathione). Purified protein sample was quantified and stored in the storage buffer as described above in the previous section.

Acetylation mimicking version of tropomyosin (ASCdc8) was expressed in BL21-DE3 and protein was purified by boiling and precipitation method as described earlier (Palani et al., 2019; Skoumpla et al., 2007). Purified tropomyosin was dialyzed against the storage buffer (50 mM NaCl, 10 mM imidazole, pH 7.5, and 1 mM DTT), flash frozen in liquid N2 and stored at −80□°C. SNAP labelling (SNAP-Surface^®^ 549, S9112S, NEB) of capping protein-beta and Rng2 1-189 was performed as per the manufactures protocol.

### SDS-PAGE and co-sedimentation assay

Purity of protein constructs was checked by running them on a 12% SDS-PAGE gel followed by staining with Coomassie blue (SimplyBlueStain, Invitrogen) and imaging on a ChemiDoc MP (BioRad).

Co-sedimentation assays were performed at 25°C by mixing 3 μM actin with varying concentrations of SpRng2(1-189), 20 min incubation followed high speed centrifugation at 100,000 g for 20 min. Equal volumes of supernatant and pellet were separated by 12% SDS-PAGE gel and stained with Coomassie blue and imaged on a ChemiDoc MP.

### Mammalian expression and imaging

*S pombe* Rng2 fragments were cloned into pCDNA3.1-eGFP using the Gibson cloning method. HEK293T and RPE-1 cells were transiently transfected with pCDNA3 containing SpRng2(1-189), SpRng2(1-189)-CaaX, SpRng2(144-189) or SpRng2(144-189)-CaaX using Lipofectamine 2000 (cat. no. 11668019, Life Technologies) following manufacturer’s instructions. Cells were transfected at ∼70% confluency for 24 h before the experiments.

For confocal microscopy imaging, 500,000 cells were transfected with 0.5 - 1 μg of DNA, and eventually 0.5 μg of pTK93 Lifeact-mCherry (#46357, Addgene). Cells were seeded and imaged on μ-Dish 35 mm (cat. no. 81156, IBIDI). Before imaging, the culture medium was replaced with phenol red–free DMEM (Opti-MEM, cat. no. 31985062, Life Technologies). Images were taken using spinning disk microscope with a 100x Apo objective, NA 1.4 (Nikon Ti-E microscope equipped with Yokogawa spinning disk unit CSU-X and an Andor iXon 897 camera).

For lattice light sheet microscopy, 1 M cells were seeded in 6 well plates containing 5 mm cover glasses (#1.5 thickness) and transfected with either 0.1 μg pCDNA3.1-eGFP-SpRng2(1-189) or with 0.5 μg pEGFP-IQGAP1 (# 30112, Addgene) and 0.5 μg pTK93 Lifeact-mCherry. Cells were imaged 16-22 hours post transfection. Cover glasses were mounted on the imaging chamber and DMEM medium was replaced by pre-warmed L-15 imaging medium (cat. no. 11415049, Gibco, Fisher scientific). Imaging was done at 37°C on a 3i second generation lattice-light-sheet microscope with a 0.71 NA LWD WI objective for excitation and a 1.1 NA WI objective for imaging and equipped with 2 Hamamatsu ORCA-Flash 4.0v3 sCMOS cameras for simultaneous dual color imaging providing a 62.5x magnification with 230×230×370 nm (x-y-z) resolution using 488 nm and 561 nm lasers for excitation. 3D volumes were recorded with 0.57 μm step size for 150 planes with 100 ms exposure.

### In vitro assay and Total Internal Reflection Fluorescence (TIRF) microscopy

#### Supported Lipid Bilayer and Experimental Chamber Preparation

The sample preparation, experimental conditions and lipid composition were similar to the ones described in previous work [Koester et al, 2016]. Glass coverslips (#1.5 borosilicate, Menzel, cat. no. 11348503, Fisher Scientific) for SLB formation were cleaned with Hellmanex III (Hellma Analytics, cat. No. Z805939, Merck) following the manufacturer’s instructions followed by thorough rinses with EtOH and MilliQ water and blow dried with N2 gas. For the experimental chamber, 0.2 ml PCR tubes (cat. no. I1402-8100, Starlab) were cut to remove the lid and conical bottom part. The remaining ring was stuck to the cleaned glass using UV glue (cat. no. NOA88, Norland Products) and three minutes curing by intense UV light at 265 nm (UV Stratalinker 2400, Stratagene). Freshly cleaned and assembled chambers were directly used for experiments.

Supported lipid bilayers (SLB) containing 98% DOPC (cat. no. 850375, Avanti Polar Lipids) and 2% DGS-NTA(Ni2+) (cat. no. 790404, Avanti Polar Lipids) lipids were formed by fusion of small uni-lamellar vesicles (SUV) that were prepared by lipid extrusion using a membrane with 100 nm pore size (cat. no. 610000, Avanti Polar Lipids). SLBs were formed by addition of 10 μl of SUV mix (at 4 mM lipid concentration) to chambers filled with 90 μl KMEH (50 mM KCl, 2 mM MgCl_2_, 1 mM EGTA, 20 mM HEPES, pH 7.2) and incubation for 30 min. Prior to addition of other proteins, the SLBs were washed 10 times by buffer exchange (always leaving 20 μl on top of the SLB to avoid damage by drying). We tested the formation of lipid bilayers and the mobility of lipids in control samples by following the recovery of fluorescence signal after photobleaching of hexa-histidine tagged GFP (His_6_-GFP) as described in (Köster et al., 2016).

#### Actin filament polymerization and tethering to SLBs

Actin was purified from muscle acetone powder form rabbit (cat. no. M6890, Merck) and labelled with Alexa488-maleimide (cat. no. A10254, Thermo Fisher) following standard protocols (Köster et al., 2016; Pardee & Spudich, 1982).

In a typical experiment, actin filaments were polymerized in parallel to SLB formation to ensure that all components of the experiment were freshly assembled before starting imaging. First 10%_vol_ of 10x ME buffer (100 mM MgCl_2_, 20 mM EGTA, pH 7.2) were mixed with unlabeled and labeled G-actin (to a final label ratio of 20%), optionally supplemented with labelled capping protein in G-actin buffer (1 mM CaCl_2_, 0.2mM ATP, 2mM Tris, 0.5 mM TCEP-HCl, pH 7.2) to a final G-actin concentration of 10 μM and incubated for 2 min to replace G-actin bound Ca^2+^ ions with Mg^2+^ ions. Polymerization of actin filaments was induced by addition of an equal amount of 2x KMEH buffer supplemented with 2 mM Mg-ATP bringing the G-actin concentration to 5 μM. After 30 min incubation time, actin filaments were added to the SLBs using blunt-cut pipette tips at a corresponding G-actin concentration of 100 nM (to ensure a homogenous mix of actin filaments, 2 μl of actin filament solution was mixed in 18 μl KMEH and then added to the SLB containing 80 μl KMEH). After 10 min of incubation, His_6_-Curly or other variants of histidine-tagged actin binding proteins at a final concentration of 10 nM were added and a short time after (1 - 5 min) binding of actin to the SLB could be observed using TIRF microscopy.

In experiments with formin, the SLB was first incubated with 10 nM His_6_-SpCdc12(FH1-FH2) and 10 nM His_6_-Curly for 20 min, then washed twice with KMEH. During the incubation time, 10%_vol_ of 10x ME buffer was mixed with unlabeled and labeled G-actin at 4 μM (final label ratio of 20%) together with 5 μM profilin and incubated for 5 min prior to addition to the SLB and imaging with TIRF microscopy.

In experiments with tropomyosin or fimbrin, actin filaments (C_G-actin_ = 1 μM) were incubated with tropomyosin at a 1:3 protein concentration ratio or with fimbrin at a 3:2 protein concentration ratio for 15 min prior to addition to the SLB (Palani et al., 2019).

In experiments with muscle myosin II filaments, we prepared muscle myosin II filaments by diluting the stock of muscle myosin II proteins (rabbit, m. psoas, cat. no. 8326-01, Hypermol) (C_myoII_ = 20 μM; 500mM KCl, 1mM EDTA, 1 mM DTT, 10 mM HEPES, pH 7.0) 10-times with MilliQ water to drop the KCl concentration to 50 mM and incubated for 5 min to ensure myosin filament formation. Myosin II filaments were further diluted in KMEH to 200 nM and added to the actin filaments bound to the SLB by His_6_-Curly by replacing 1/10 of the sample buffer with the myosin II filament solution and supplemented with 0.1 mM Mg-ATP as well as a mix of 1 mM Trolox (cat. no. 648471, Merck), 2 mM protocatechuic acid (cat. no. 03930590, Merck) and 0.1 μM protocatechuate 3,4-dioxygenase (cat. no. P8279, Merck) to minimize photobleaching. To summarize, the final buffer composition was 50mM KCl, 2mM MgCl_2_, 1mM EGTA, 20mM HEPES, 0.1mM ATP, 1 mM Trolox, 2 mM protocatechuic acid and 0.1 μM protocatechuate 3,4-dioxygenase at pH 7.2 containing actin filaments (C_G-actin_ = 100 nM) and myosin II filaments (C_myoII_ = 20 nM). It was important to keep the pH at 7.2, as changes in pH would affect motor activity. As reported earlier, myosin filaments started to show actin network remodeling activity after about 10-15 min of incubation (Köster et al., 2016; Mosby et al., 2020).

#### TIRF microscopy

Images were acquired using a Nikon Eclipse Ti-E/B microscope equipped with perfect focus system, a Ti-E TIRF illuminator (CW laser lines: 488nm, 561nm and 640nm) and a Zyla sCMOS 4.2 camera (Andor, Oxford Instruments, UK) controlled by Andor iQ3 software (https://andor.oxinst.com/products/iq-live-cell-imaging-software/).

### Image analysis

Images were analyzed using ImageJ (http://imagej.nih.gov/ij).

Curvature was measured by fitting ellipses to match the actin filament contour by hand, while measuring first fully formed rings before curved actin filament segments and by going from the highest curvatures down to lower curvatures in each image with a cut off for measurements at curvatures smaller than 0.1 μm^−1^ or at 30-40 measurements per image (see examples in Figure 1 – figure supplement 1D; Figure 1-figure supplement 2B).

The actin ring contraction rate upon myosin II filament action was measured by generating kymographs based on a line (3 pixels width) dividing the ring into two equal halves.

The 3D projection animation of HEK293 cells expressing EGFP-IQGAP1 and mRuby-LifeAct was generated with the 3Dscript plugin for ImageJ (Schmid et al., 2019).

### Data plotting and statistics

Graphs were generated using OriginPro (version 2019b, OriginLab, USA). All box plots depict individual data points, mean (circle), median (central line) and standard deviation (top and bottom lines).

**Table 1:** Summary of experimental results.

| Experiment | Curvature C ($\mu\text{m}^{-1}$) | Rings / FoV | p Value (C = C<sub>His6-Rng2(1-189)</sub>) |
| --- | --- | --- | --- |
| SLB + His<sub>6</sub>-Rng2(1-189) | 1.7 (± 0.5) | 21 (± 7) | 1 |
| glass + His<sub>6</sub>-Rng2(1-189) | 0.6 (± 0.3) | 0.5 (± 0.7) | 0.003 |
| SLB + His<sub>6</sub>-Rng2(150-250) | 0.3 (± 0.1) | 1.1 (± 0.9) | 0.0002 |
| SLB + His<sub>6</sub>-Rng2(1-189)$\Delta$(154-169) | 0.6 (± 0.2) | 0.1 (± 0.4) | 0.0006 |
| SLB + His<sub>6</sub>-Rng2(41-189) | 0.6 (± 0.2) | 0.4 (± 0.5) | 0.0007 |
| SLB + His<sub>6</sub>-Rng2(1-147) | 0.7 (± 0.2) | 0 | 0.003 |
| SLB + Rng2(1-189)-His<sub>6</sub> | 1.4 (± 0.4) | 24 (± 6) | 0.08 |
| SLB + His<sub>6</sub>-Rng2(1-189) + His<sub>6</sub>-Formin | 1.7 (± 0.4) | 76 (± 34) | 0.7 |
| SLB + His<sub>6</sub>-Rng2(1-189) + Tropomyosin-actin | 1.4 (± 0.6) | 33 (± 7) | 0.3 |
| SLB + His<sub>6</sub>-Rng2(1-189) + Fimbrin-actin | 0.6 (± 0.4) | 3 (± 2) | 0.0005 |
| SLB + His<sub>6</sub>-Rng2(1-189) + Myosin II | 2.8 (± 0.7) | 95 (± 18) | 0.0006 |
| SLB + His<sub>6</sub>-Iqg1(1-330) (S.C.) | 1.1 (± 0.4) | 9 (± 6) | 0.03 |
| SLB + His<sub>6</sub>-IQGAP1(1-678) (H.S.) | 1.0 (± 0.2) | 13 (± 5) | 0.005 |
| HEK293T + EGFP-Rng2(1-189) | 2.3 (± 0.4) | 5 (± 3) | N.A. |
| RPE-1 + EGFP-Rng2(1-189) | 1.9 (± 0.6) | 12 (± 8) | N.A. |
| SLB + α-actinin-His<sub>6</sub> | 0.3 (± 0.1) | | N.A. |
| SLB + His<sub>10</sub>-SNAP-EzrinABD | 0.5 (± 0.3) | | N.A. |

**Table 2:**
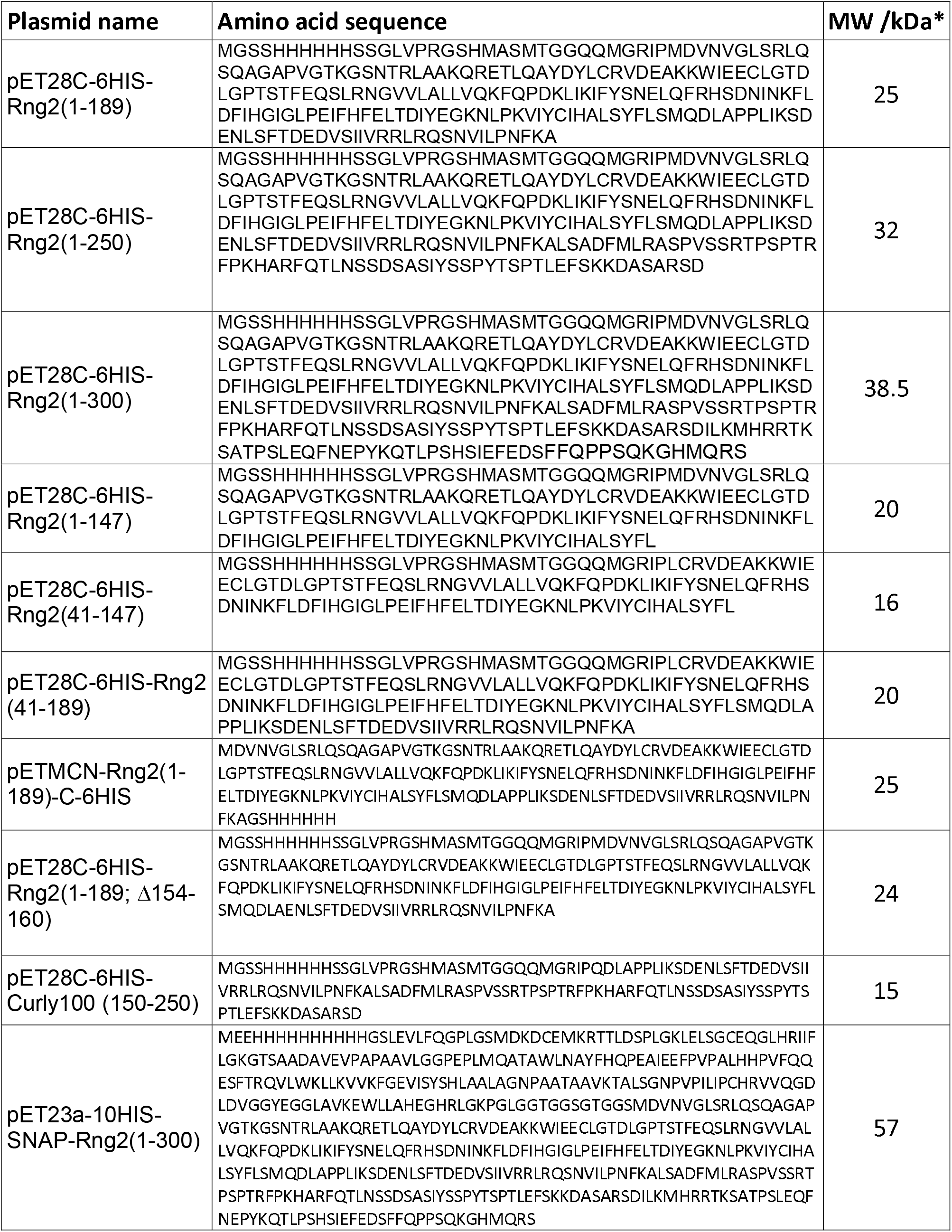
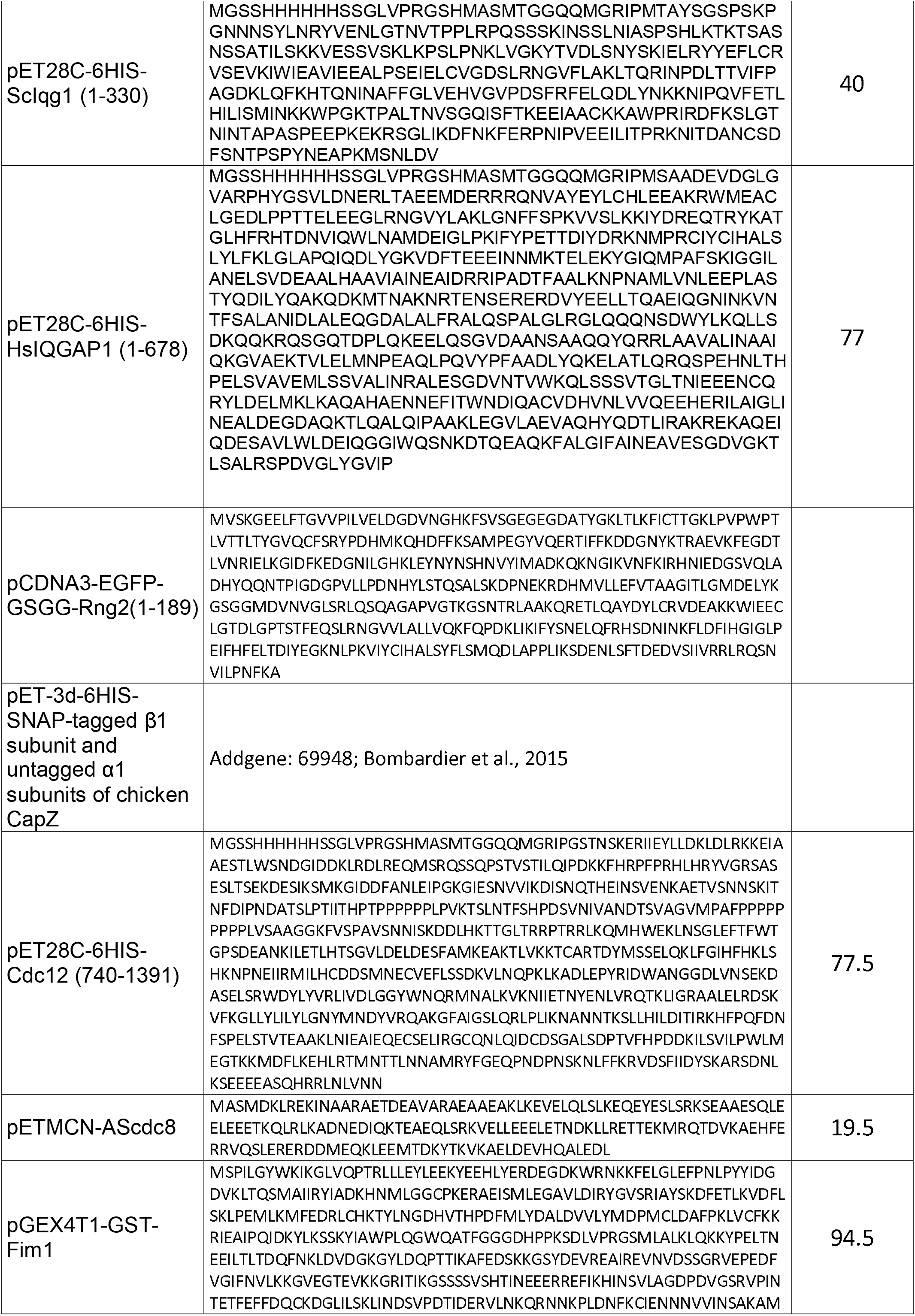
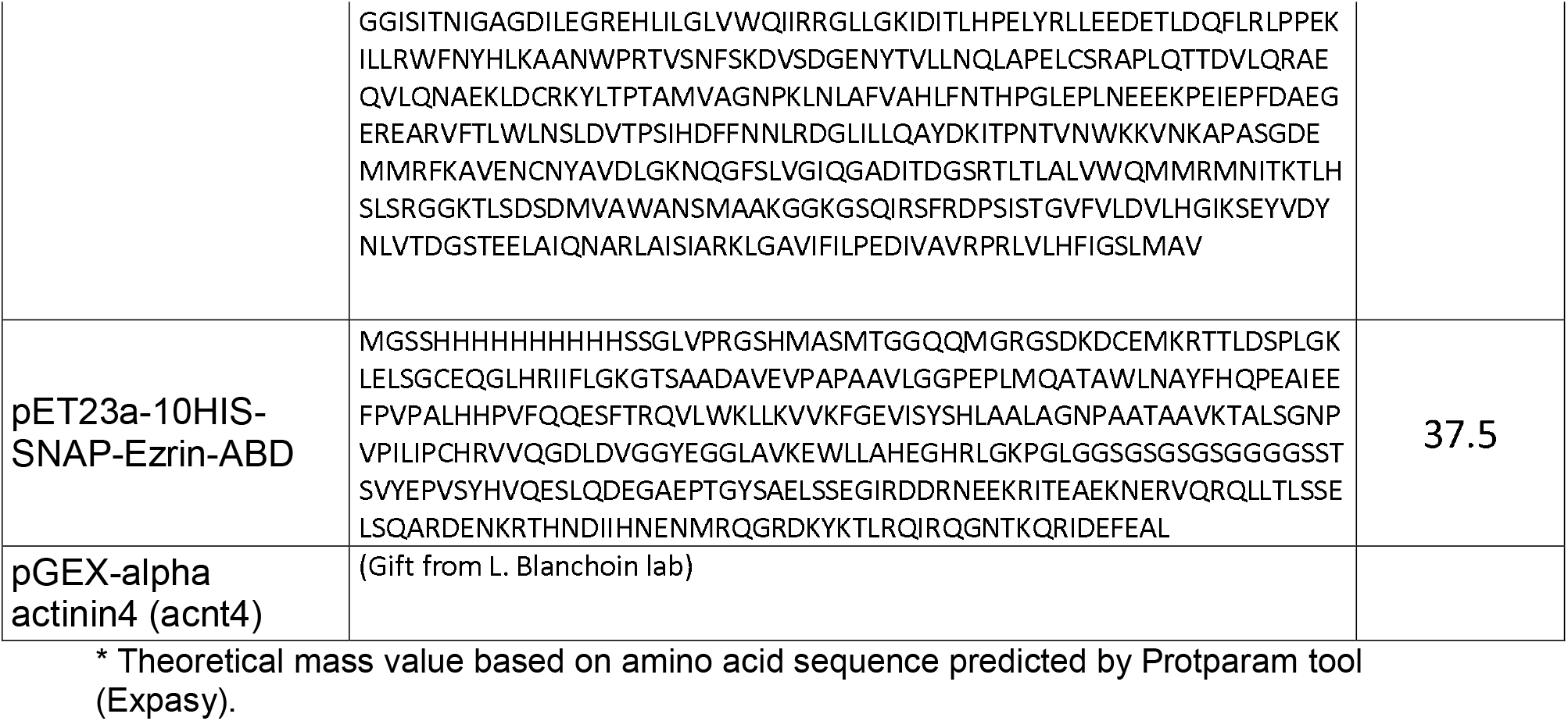
Plasmids used in this study.

## Acknowledgement

The authors would like to thank Dr. Gayathri Panaghat (IISER Pune, India), Dr. Minhaj Sirajuddin (InStem, Bangalore, India), for insightful discussions and Prof. Gijsje Koenderink (TU Delft, Netherlands) and Prof. Rob Cross (University of Warwick, UK) for helpful comments on the manuscript. The work was supported by a Wellcome Investigator Award (WT 101885MA) and an ERC advanced grant (ERC-2014-ADG N° 671083) to MKB. DVK thanks the Wellcome-Warwick Quantitative Biomedicine Programme for funding (Wellcome Trust ISSF, RMRCB0058). SP thanks the DBT-IISc partnership funds and IISc start-up grant. SG was supported by an international Chancellor’s fellowship of the University of Warwick. SC was supported by a research development grant of the University of Warwick. EIM was supported by the A*STAR Research Attachment Programme (ARAP) PhD studentship.

## Competing Interests

The authors have no competing interests to declare.

## Author contributions

SP: Conceptualization, Funding acquisition, Data curation, Formal analysis (protein structure), Investigation, Methodology, Project administration, Writing-original draft, Writing-review and editing;

AS: Investigation (protein purification and sedimentation assay);

SG: investigation (transfection and culture of mammalian cells);

SC: investigation (cell imaging using lattice-light-sheet microscopy);

EIM: Investigation (transfection and culture of mammalian cells);

MBK: Conceptualization, Supervising, Funding acquisition, Methodology, Project administration, Writing-original draft, Writing-review and editing;

DVK: Conceptualization, Supervising, Funding acquisition, Data curation, Formal analysis, Investigation, Methodology, Project administration, Writing-original draft, Writing-review and editing.

---

**Figure 1 – figure supplement 1.**
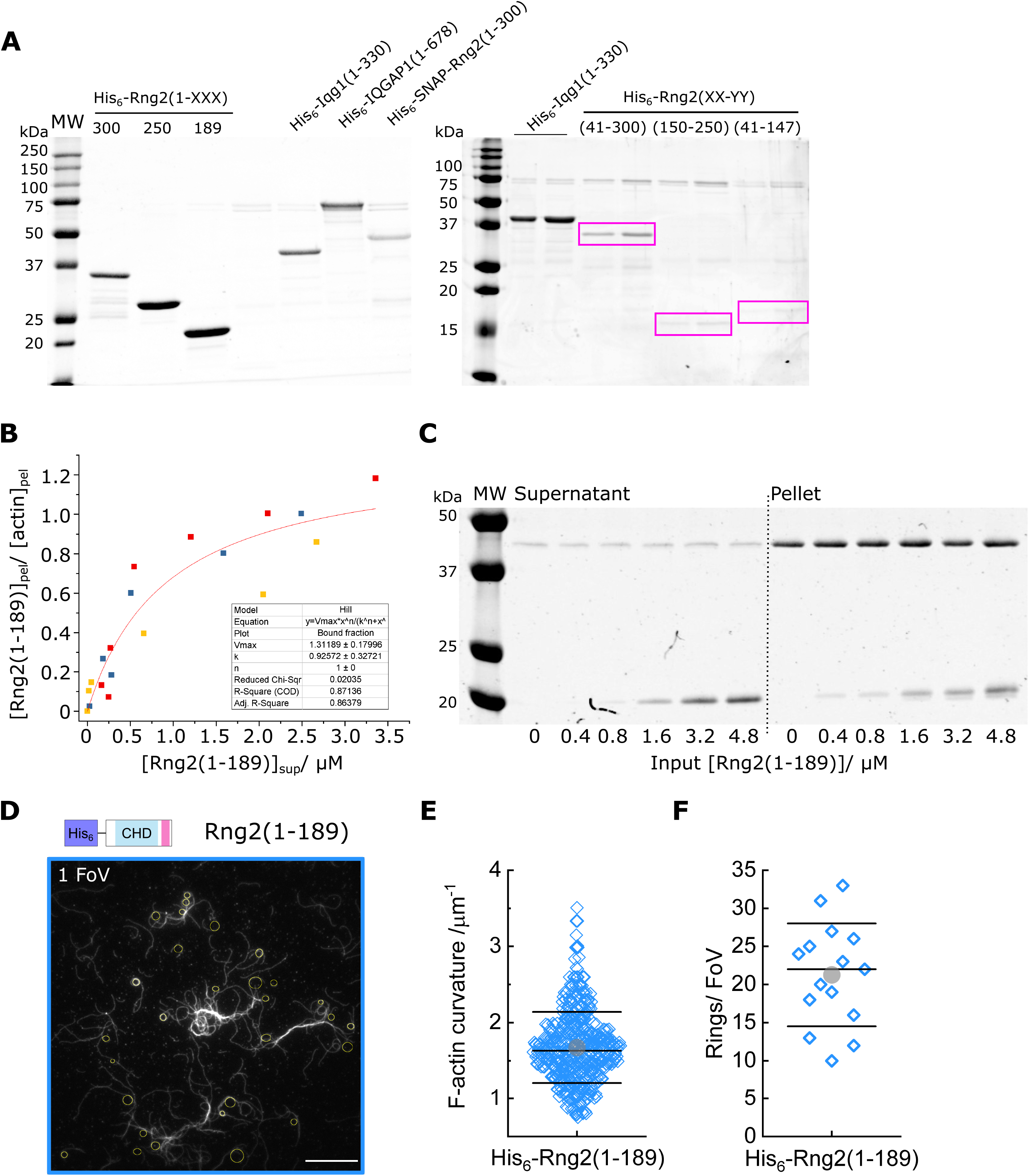
(A) Coomassie stained SDS-PAGE of the different constructs used in this study; purple boxes highlight the position of the relevant protein band. (B) Quantification of Rng2(1-189) found in the pellet and supernatant fractions after co-sedimentation with actin filaments; the solid line depicts a fit with the hill-function for non-cooperative binding (n = 1) and K_d_ = 0.9 (± 0.3) μM-1. (C) Example of a Coomassie stained SDS-PAGE showing the supernatant and pellet fractions of different Rng2(1-189) concentrations after co-sedimentation with 3 μM actin. (D) TIRF microscopy image (1 FoV) of actin filaments (Alexa488) bound to SLB tethered His_6_-curly; circles show curvature measurements; scale bar 10 μm. (E) Box plot of actin filament curvatures induced by SLB tethered His_6_-curly (same as in Fig. 1 C but with linear scale). (F) Box plot depicting the number of full actin rings per field of view (FoV) induced by SLB tethered His_6_-Rng2(1-189); N = 15 FoVs from three independent samples.

**Figure 1-figure supplement 2.**
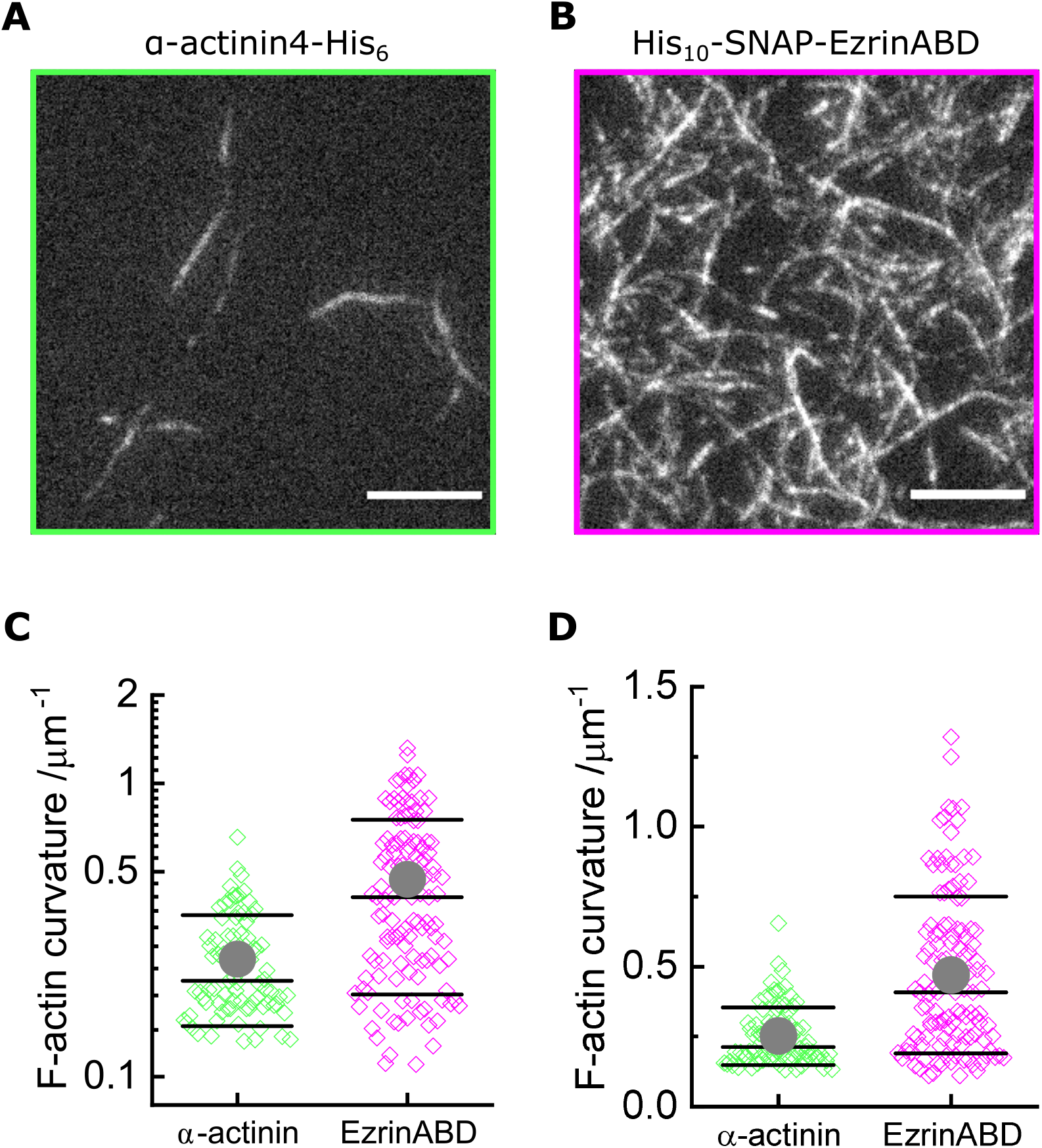
(A) TIRF microscopy image of actin filaments (Alexa488, C_actin_ = 100 nM) bound to SLB tethered α-actinin-His_6_ (Cα-act = 10 nM); scale bar 5 μm. (B) TIRF microscopy image of actin filaments (Alexa488, C_actin_ = 100 nM) bound to SLB tethered His_10_-EzrinABD (C_EzrABD_ = 10 nM); scale bar 5 μm. (C) Curvature measurements of actin filament rings and curved segments; α-actinin-His_6_ : N = 85 obtained from 10 field of views from 4 individual experiments; His_10_-EzrinABD : N = 127 obtained from 9 field of views from 3 individual experiments. (D) Same data as (C) but in linear scale.

**Figure 2- Figure supplement 1.**
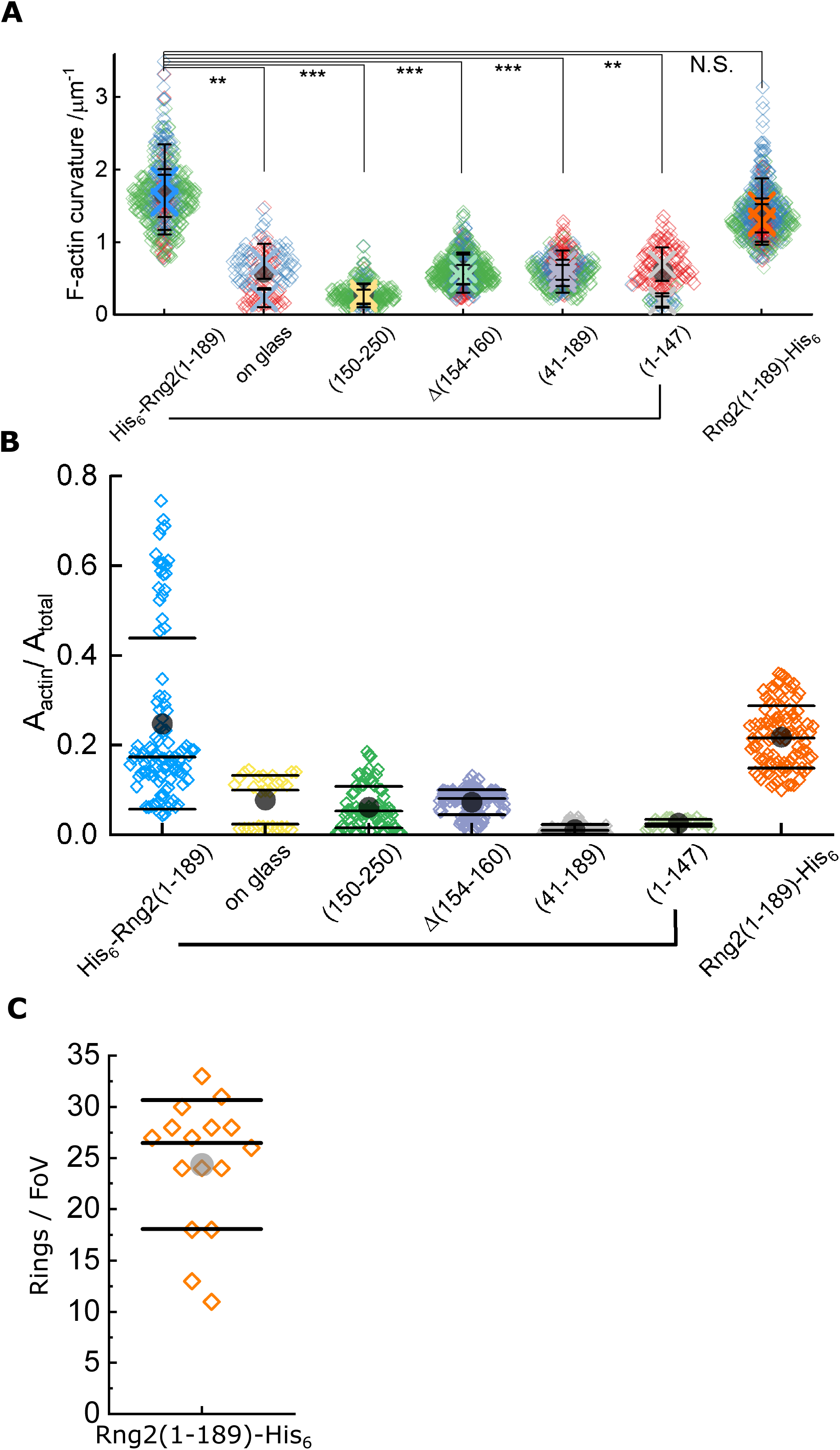
Comparison of actin binding to SLBs by curly constructs. (A) Statistical comparison of average actin filament curvatures induced by Rng2 constructs as shown in Fig. 2G; plotted are the mean value ± standard deviation of independent experiments, comparison of mean values with Anova one-way test; **: p < 0.005, ***: p < 0.001, N.S.: non-significant difference between the mean values. (B) Relative area of actin coverage of the SLB in the field of view indicating the actin binding capacity of the Rng2 constructs used in Fig. 2. Number of full actin rings per field of view induced by SLB tethered Rng2(1-189)-His_6_; N = 16 FoVs from three independent samples.

**Figure 3-figure supplement 1.**
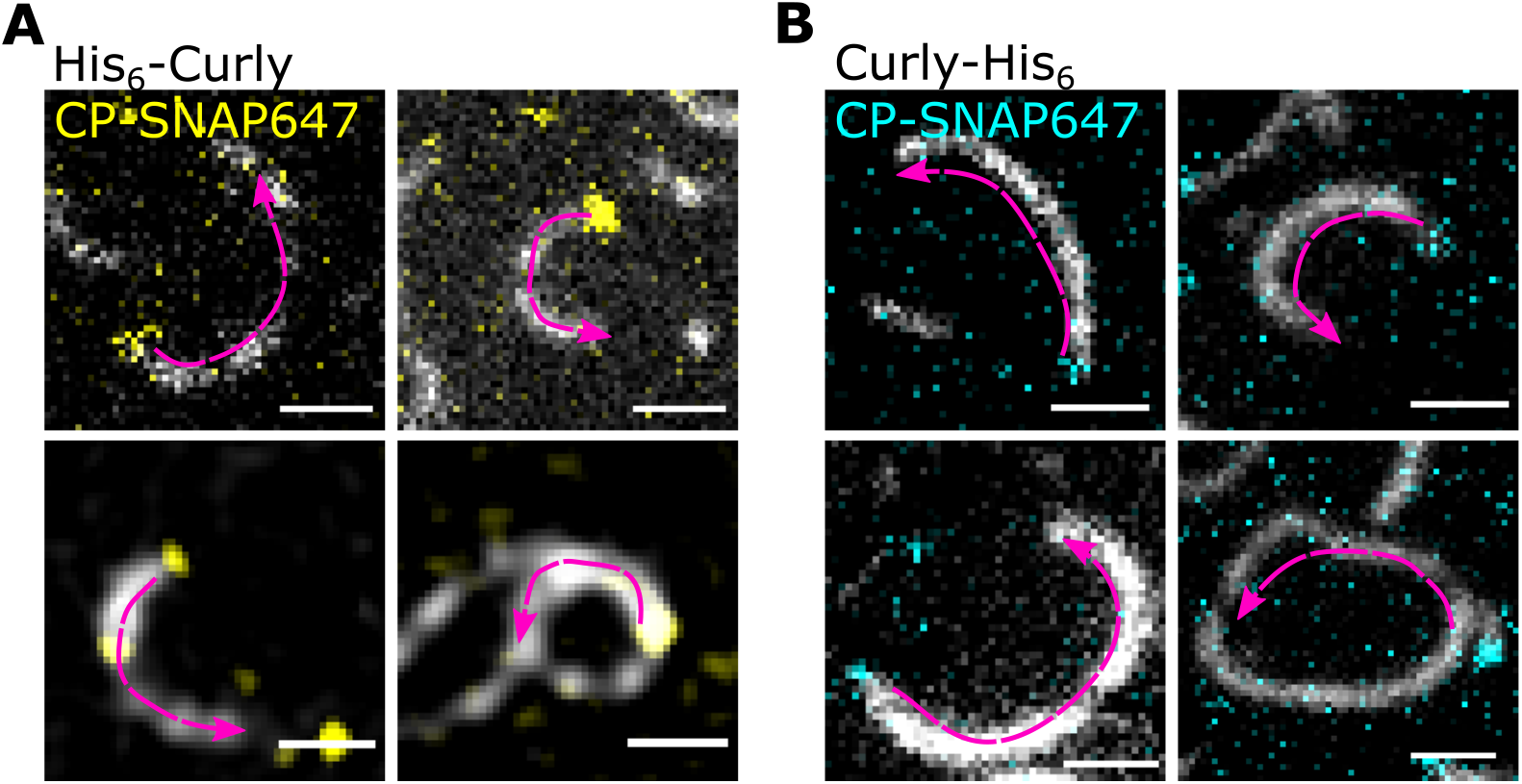
(A) TIRF microscopy images of actin filaments (Alexa488) with the plus end marked with SNAP647-tagged capping protein binding to His_6_-Rng2(1-189); scale bar: 1 μm. (B) TIRF microscopy images of actin filaments (Alexa488) with the plus end marked with SNAP647-tagged capping protein binding to Rng2(1-189)-His_6_; scale bar: 1 μm. Arrows indicate the actin bending orientation.

**Figure 3-figure supplement 2.**
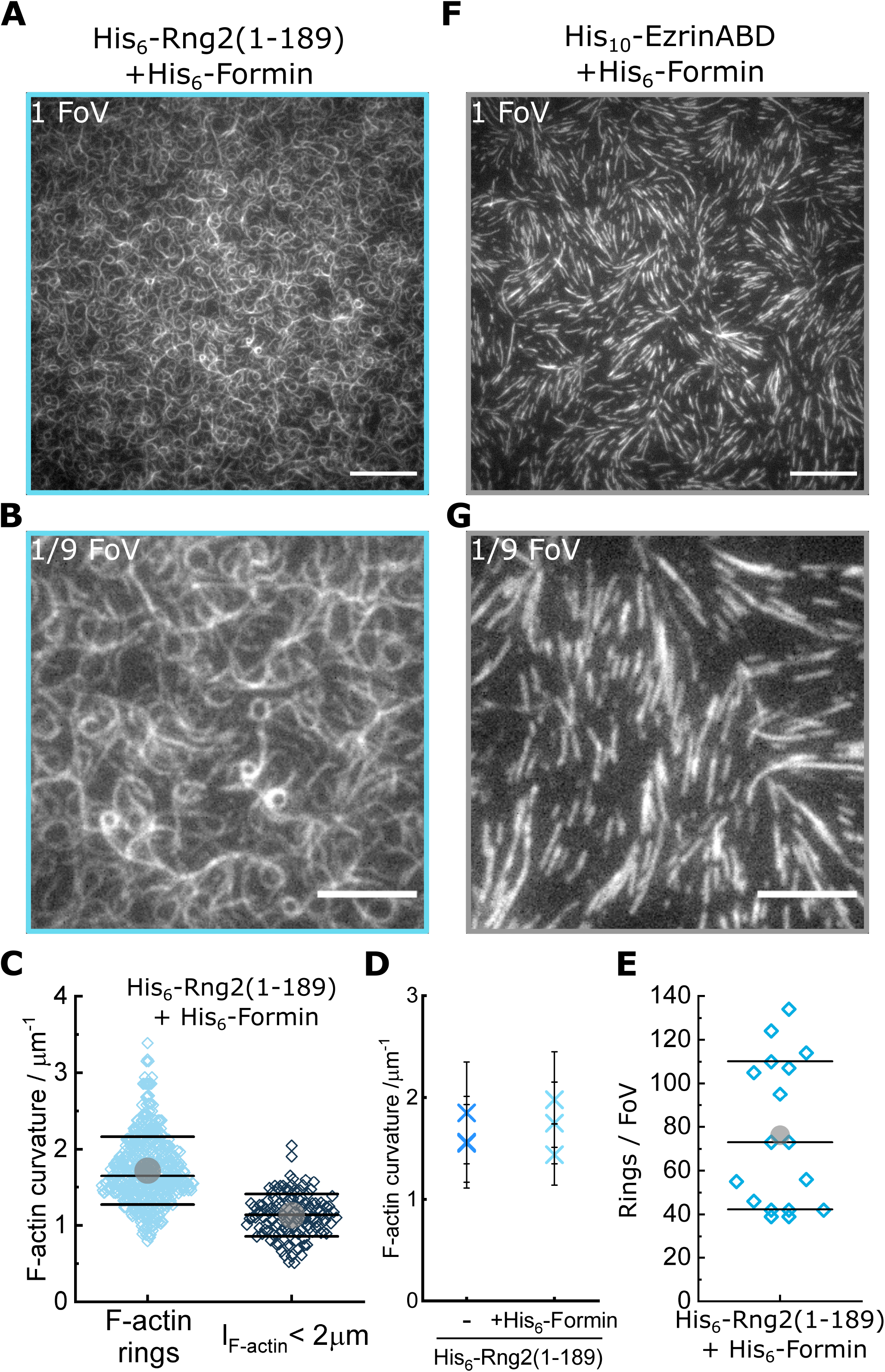
(A) TIRF microscopy image (full field of view) of actin filaments (Alexa488) polymerized by membrane tethered His_6_-formin in the presence of His_6_-Rng2(1-189); scale bar: 10 μm. (B) Zoom of (A) showing 1/9 of the field of view; scale bar: 5 μm. (C) Data from Fig. 3D plotted in linear scale. (D) Comparison of mean actin ring curvatures of independent experiments with formin induced actin polymerization on curly containing membranes (data from Fig. 3 D) and with actin filaments landing on curly decorated membranes (data from Fig. 1 C) indicating that the curly induced ring curvature is independent of the type of actin polymerization. (E) Number of full actin rings per field of view when actin filaments are polymerized by membrane tethered His_6_-formin in the presence of His_6_-Rng2(1-189); N = 17 FoVs from three independent samples. (F) TIRF microscopy image (full field of view) of actin filaments (Alexa488) polymerized by membrane tethered His_6_-formin in the presence of His_10_-EzrinABD; scale bar: 10 μm. (G) Zoom of (F) showing 1/9 of the field of view; scale bar: 5 μm.

**Figure 3-figure supplement 3.**
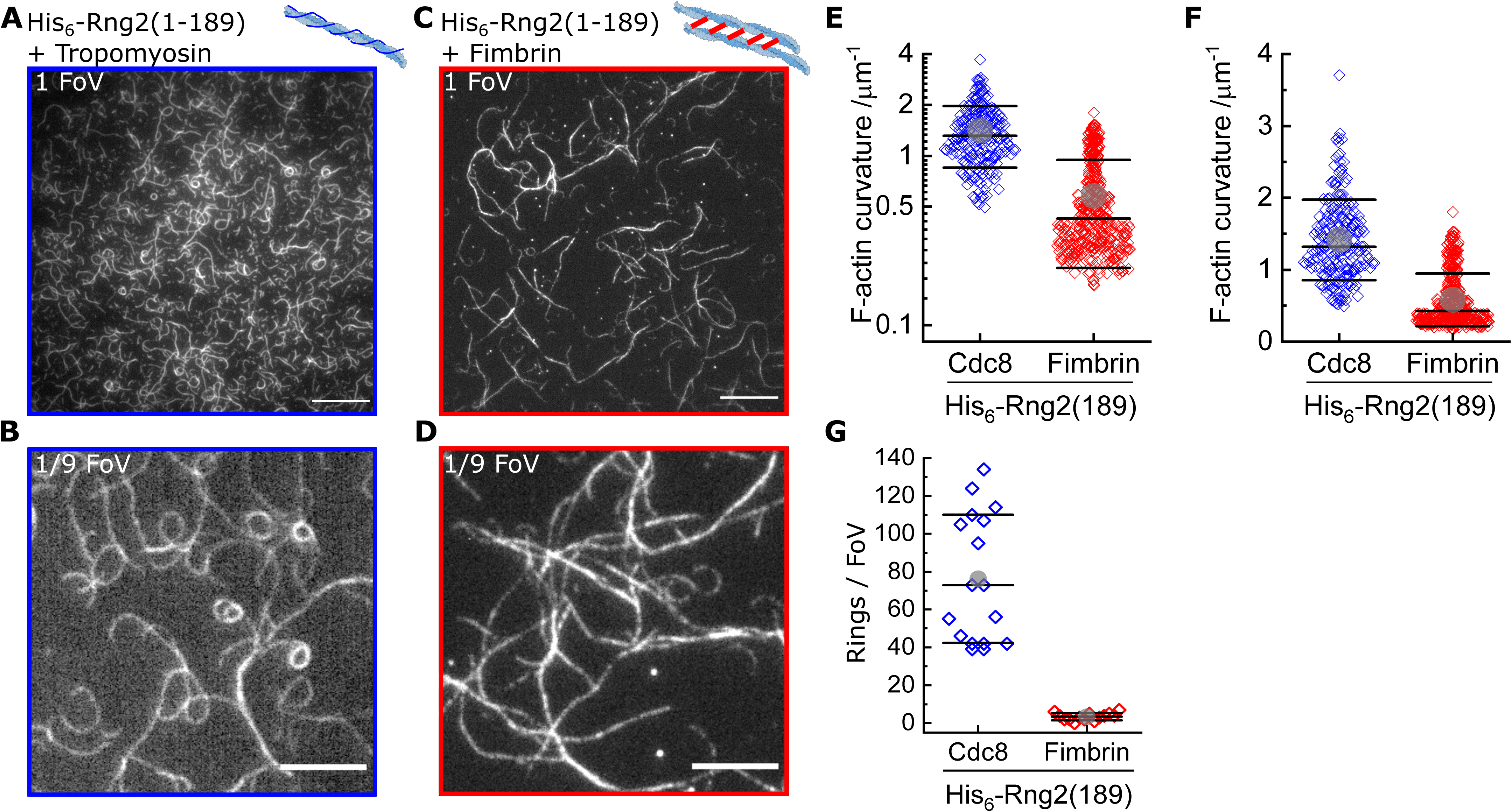
(A) TIRF microscopy image (one field of view) of actin filaments (Alexa488) pre-incubated with tropomyosin (Cdc8) bound to membrane tethered His_6_-Rng2(1-189); scale bar: 10 μm. (B) TIRF microscopy image (1/9 field of view) of actin filaments (Alexa488) pre-incubated with tropomyosin (Cdc8) bound to membrane tethered His_6_-Rng2(1-189); scale bar: 5 μm. (C) TIRF microscopy image (representing one field of view) of actin filaments (Alexa488) pre-incubated with fimbrin bound to membrane tethered His_6_-Rng2(1-189); scale bar: 10 μm. (D) TIRF microscopy image (representing 1/9 field of view) of actin filaments (Alexa488) pre-incubated with fimbrin bound to membrane tethered His_6_-Rng2(1-189); scale bar: 5 μm. (E) Curvature measurements of actin filament rings and curved segments; Cdc8 (blue): N = 204 from 9 field of views of 3 individual experiments; fimbrin (red): N = 407 from 20 field of views of 3 independent experiments. (F) Data from (E) plotted in linear scale. (G) Number of full actin rings per field of view, N(Cdc8) = 16, N(fimbrin) = 14 from three independent experiments.

**Figure 3-figure supplement 4.**
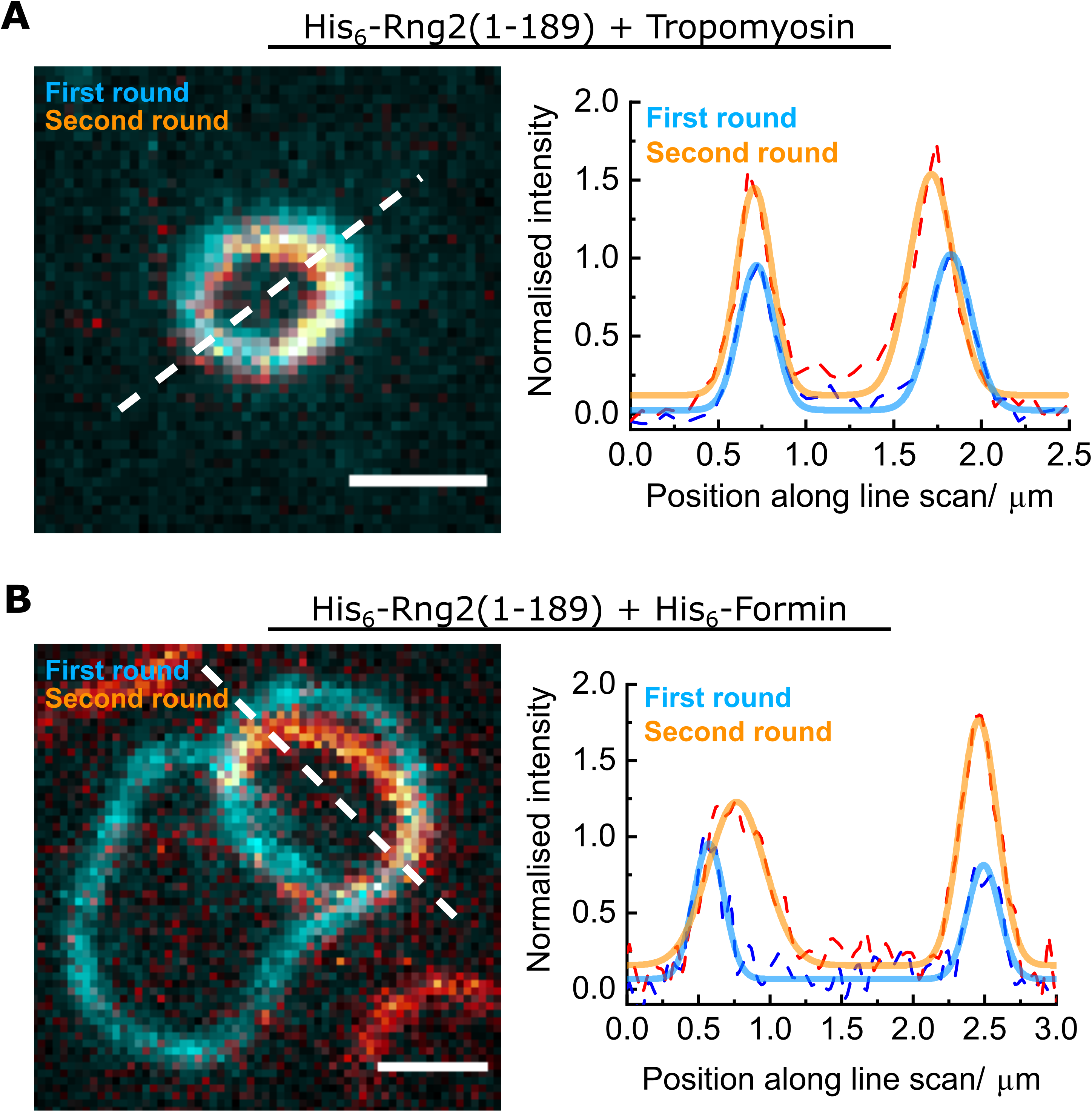
(A) Left: TIRF microscopy image overlay showing multiple ring formation of a tropomyosin coated actin filament (Alexa488) during binding to membrane tethered His_6_-Rng2(1-189); the first ring formed is colored in cyan, the second ring (in orange) was highlighted by subtracting the image of the first ring from the image stack; scale bar: 1 μm. Right: Intensity line scan (3 pixels width) along the white dashed line and corresponding Gaussian peak fits (colored dashed lines). (B) Left: TIRF microscopy image overlay showing multiple ring formation of a polymerizing actin filament (Alexa488) by membrane tethered His_6_-formin in presence of membrane tethered His_6_-Rng2(1-189); the first ring formed is colored in cyan, the second ring (in orange) was highlighted by subtracting the image of the first ring from the image stack; scale bar: 1 μm. Right: Intensity line scans (3 pixels width) along the white dashed line and corresponding Gaussian peak fits (colored dashed lines).

**Figure 3-figure supplement 5.**
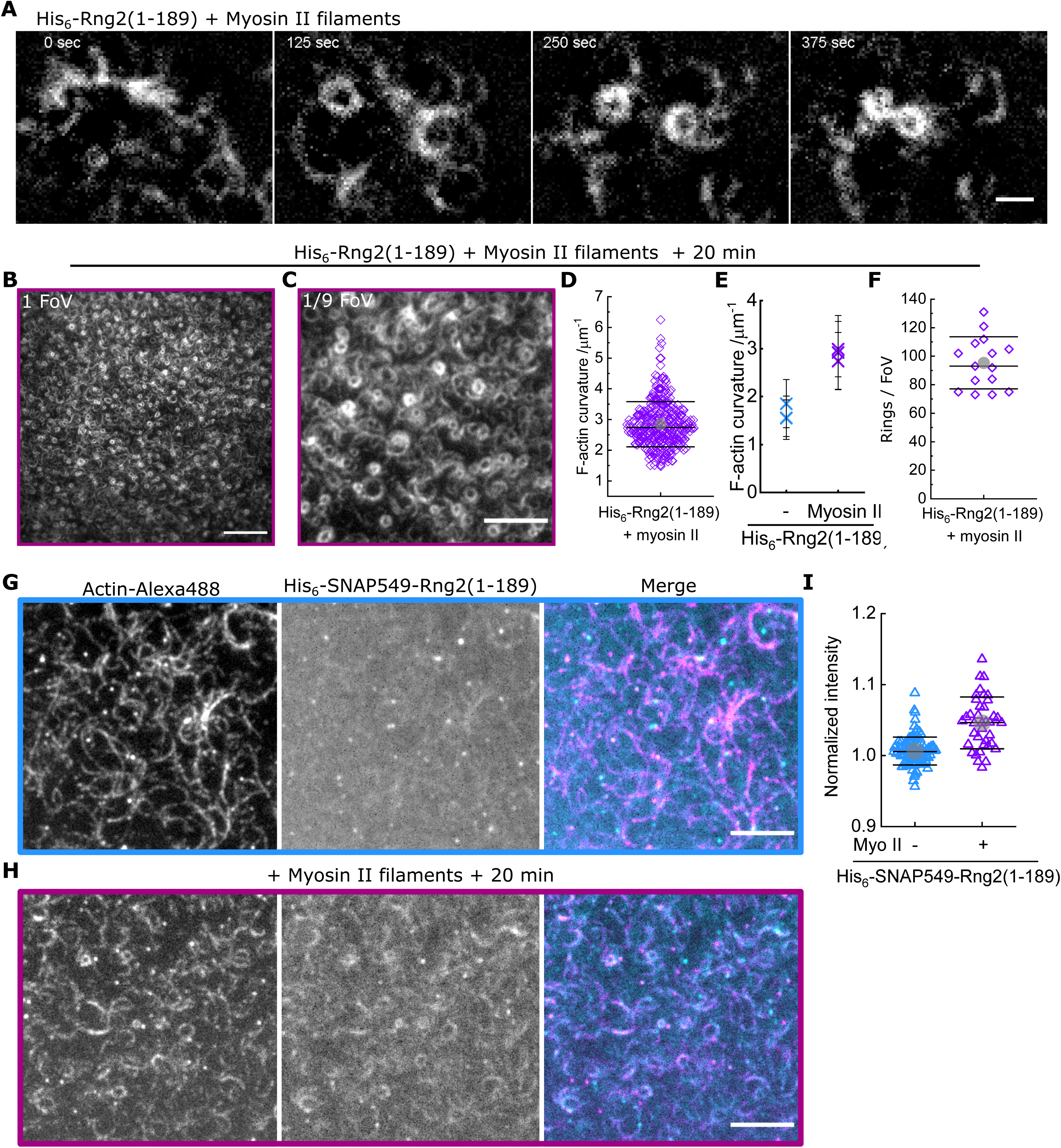
(A) TIRF microscopy image time series showing rabbit muscle myosin II filament driven ring formation, sliding and contraction of actin filaments (Alexa488) bound to membrane tethered His_6_-Rng2(1-189); scale bar: 1 μm. (B) TIRF microscopy image (representing one field of view) of actin filaments (Alexa488) 20 min after addition of rabbit muscle myosin II filaments on His_6_-curly containing SLBs; scale bar: 10 μm. (C) Zoom of (B) showing 1/9 field of view; scale bar: 5 μm. (D) Data from Fig. 3G plotted in linear scale. (E) Comparison of mean actin ring curvatures of individual experiments induced by His_6_-Rng2(1-189) alone (data from Fig. 1 C) and in presence of rabbit muscle myosin II filament after 20 min of incubation (data from Fig. 3 G). (F) Number of full actin rings per field of view of actin filaments 20 min after addition of rabbit muscle myosin II filaments on His_6_-Rng2(1-189) containing SLBs; N = 15 from three independent experiments. (G) Two color TIRF microscopy image of actin filaments (Alexa488, magenta) bound to membrane tethered fluorescently labelled His_6_-SNAP-Rng2(1-189) (Surface549, cyan) before… (H) …and 20 min after addition of rabbit muscle myosin II filaments; scale bar 5 μm. (I) Box plot of His_6_-SNAP-Rng2(1-189) (Surface549) intensities under curved actin filaments before and after addition of rabbit muscle myosin II filaments; intensities were normalized to the average intensity of the entire field of view.

**Figure 4-figure supplement 1.**
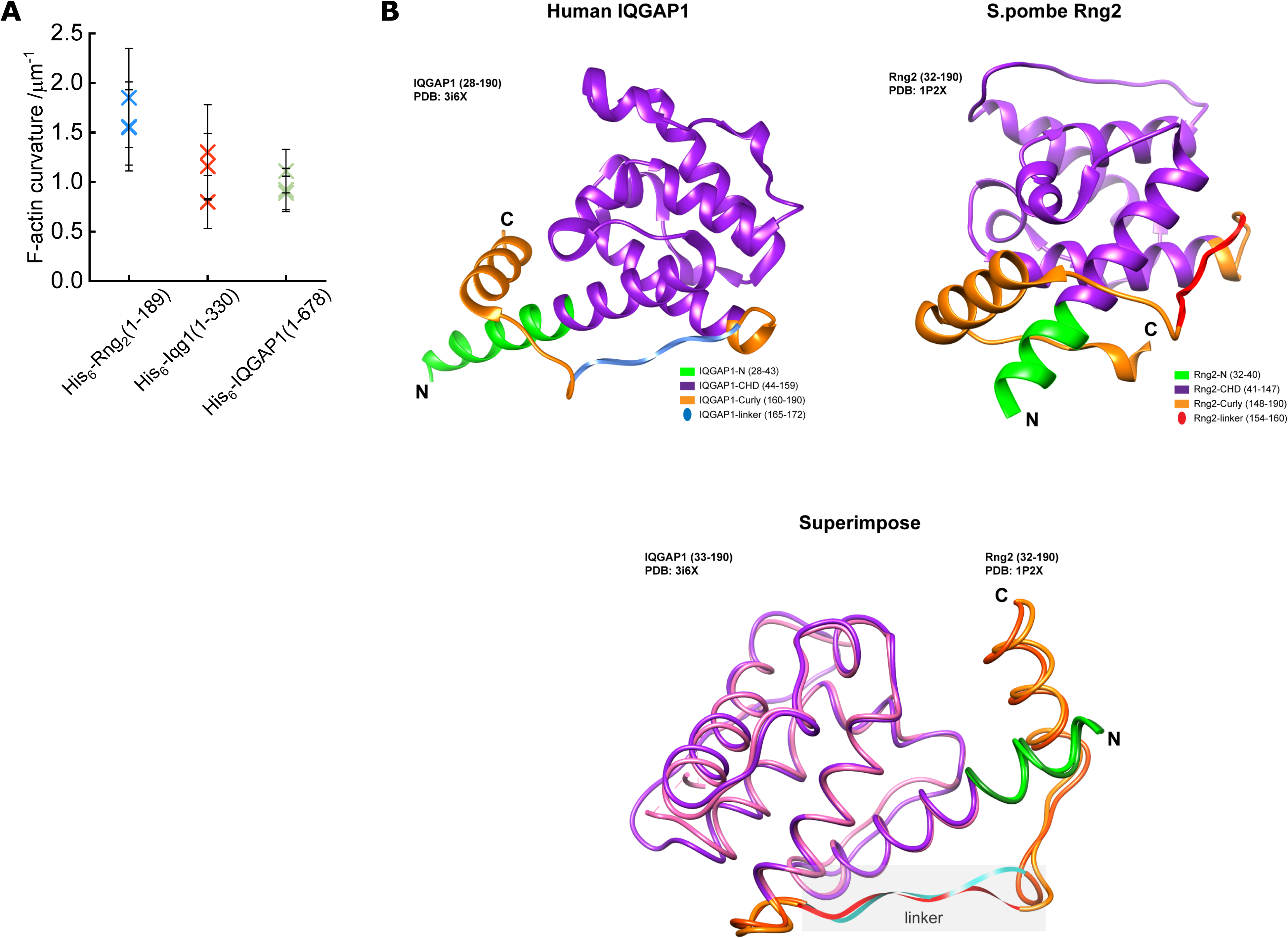
(A) Comparison of mean actin ring curvatures of individual experiments induced by His_6_-Rng2(1-189) (data from Fig. 1 C), His_6_-Iqg1(1-330) and His_6_-IQGAP1(1-678) (data from Fig. 4 C). (B) Depiction of structure predictions and overlay of H. sapiens IQGAP1(28-190) and S. pombe Rng2(32-190) indicating the strong similarity between the linker regions of both proteins that are thought to be important for actin bending.

**Figure 4-figure supplement 2.**
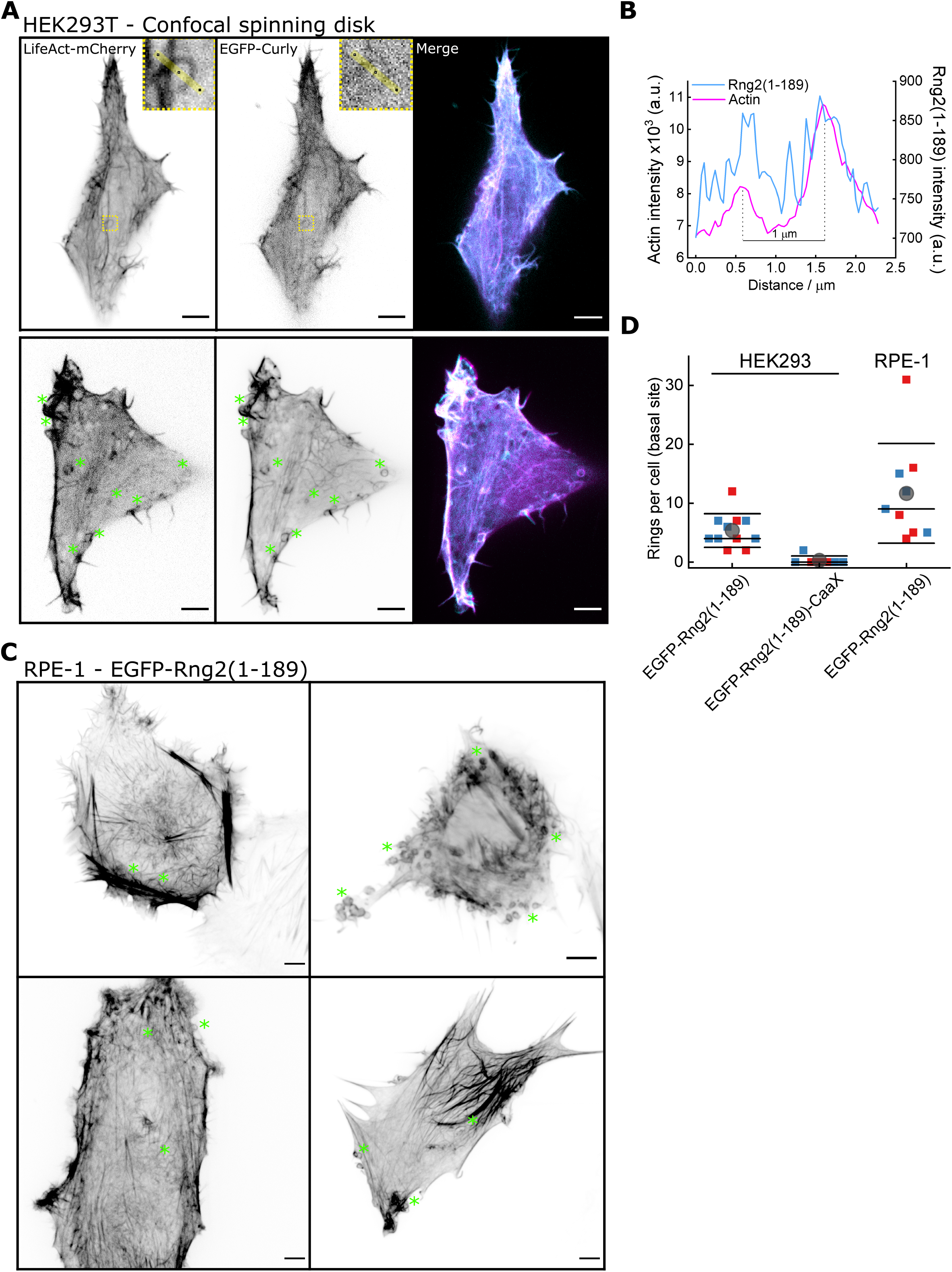
(A) Confocal microscopy images of HEK293T cells transfected with LifeAct-mCherry (magenta) and EGFP-Rng2(1-189) (cyan); inlet shows a ring structure showing both LifeAct and curly labelling; green stars indicate location of ring structures; images show maximum intensity projection of the basal cell section of ~2μm; scale bar 5 μm. (B) Line scan of the ring in (A) depicting the intensity profile of LifeAct-mCherry and EGFP-Rng2(1-189). (C) Confocal microscopy images of RPE-1 cells transfected with EGFP-Rng2(1-189); green stars indicate location of ring structures; images show maximum intensity projection of the basal cell section of ~2μm; scale bar 5 μm. (D) Quantification of actin rings found in the basal section of each cell, red and blue indicate the two independent experiments.

**Figure 4-figure supplement 3.**
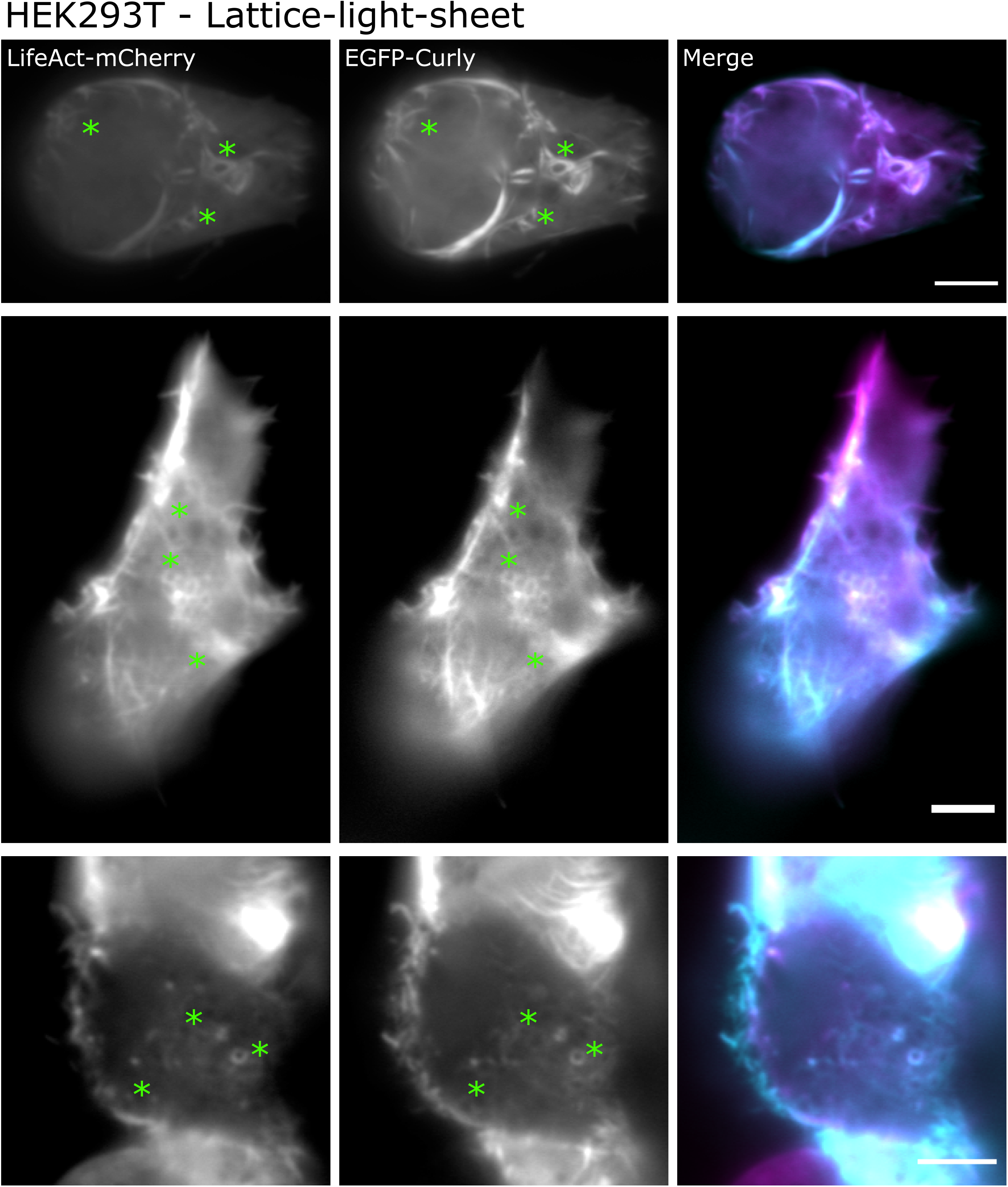
Lattice-light-sheet microscopy images (maximum intensity projections of middle or basal section) of HEK293T cells transfected with LifeAct-mCherry (magenta) and EGFP-Rng2(1-189) (cyan); green stars indicate location of ring structures; scale bar: 5 μm.

**Figure 4-figure supplement 4.**
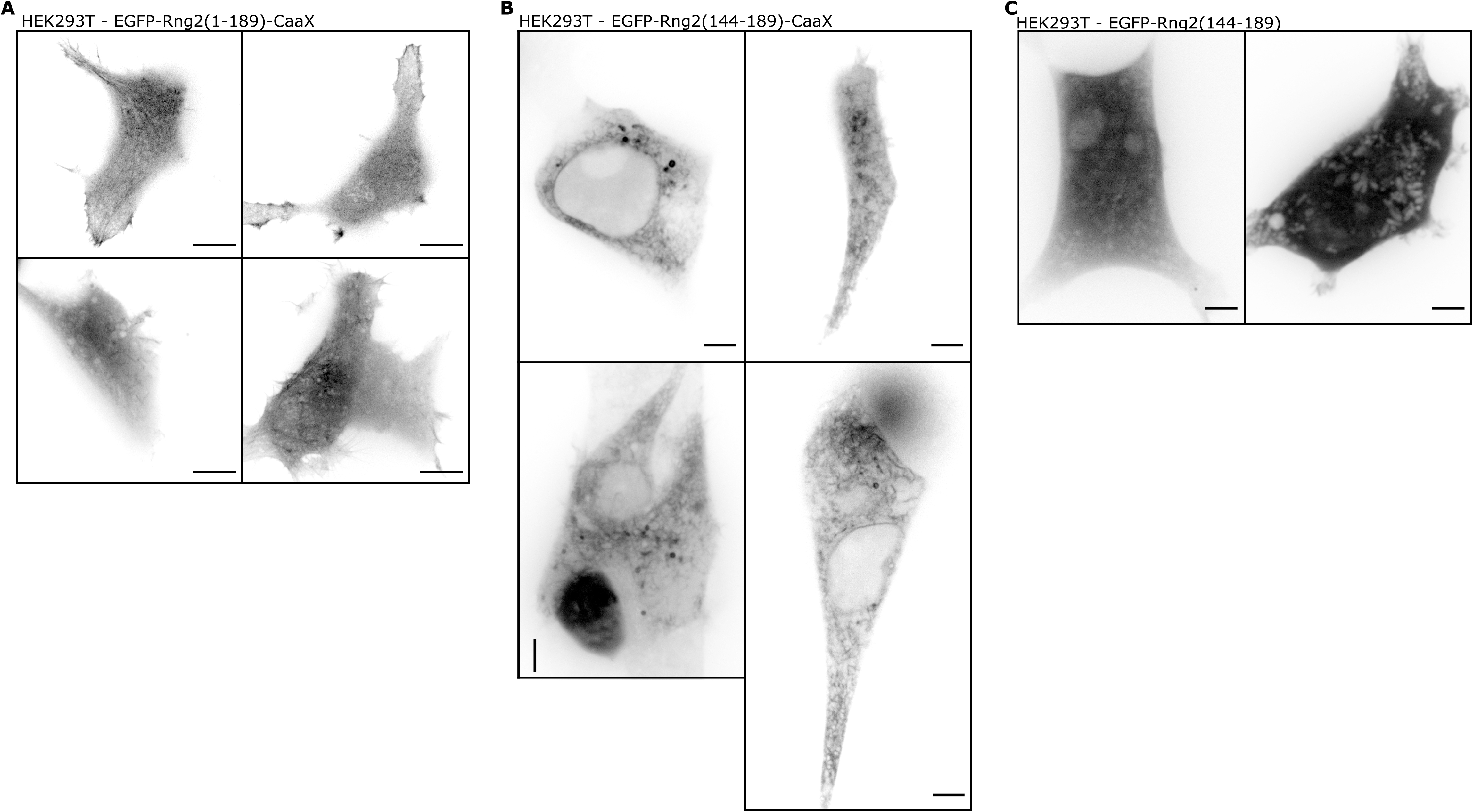
(A) Confocal microscopy images of HEK293T cells transfected with EGFPRng2(1-189)-CaaX; images show maximum intensity projection of the basal cell section (~2μm)); scale bar 5 μm. (B) Confocal microscopy images of HEK293T cells transfected with EGFP-Rng2(144-189)-CaaX; images show maximum intensity projection of the basal cell section (~2μm)); scale bar 5 μm. (C) Confocal microscopy Example images of HEK293T cells transfected with EGFP-Rng2(44-189); images show maximum intensity projection of the basal cell section (~2μm)); scale bar 5 μm.

**Figure 4-figure supplement 5.**
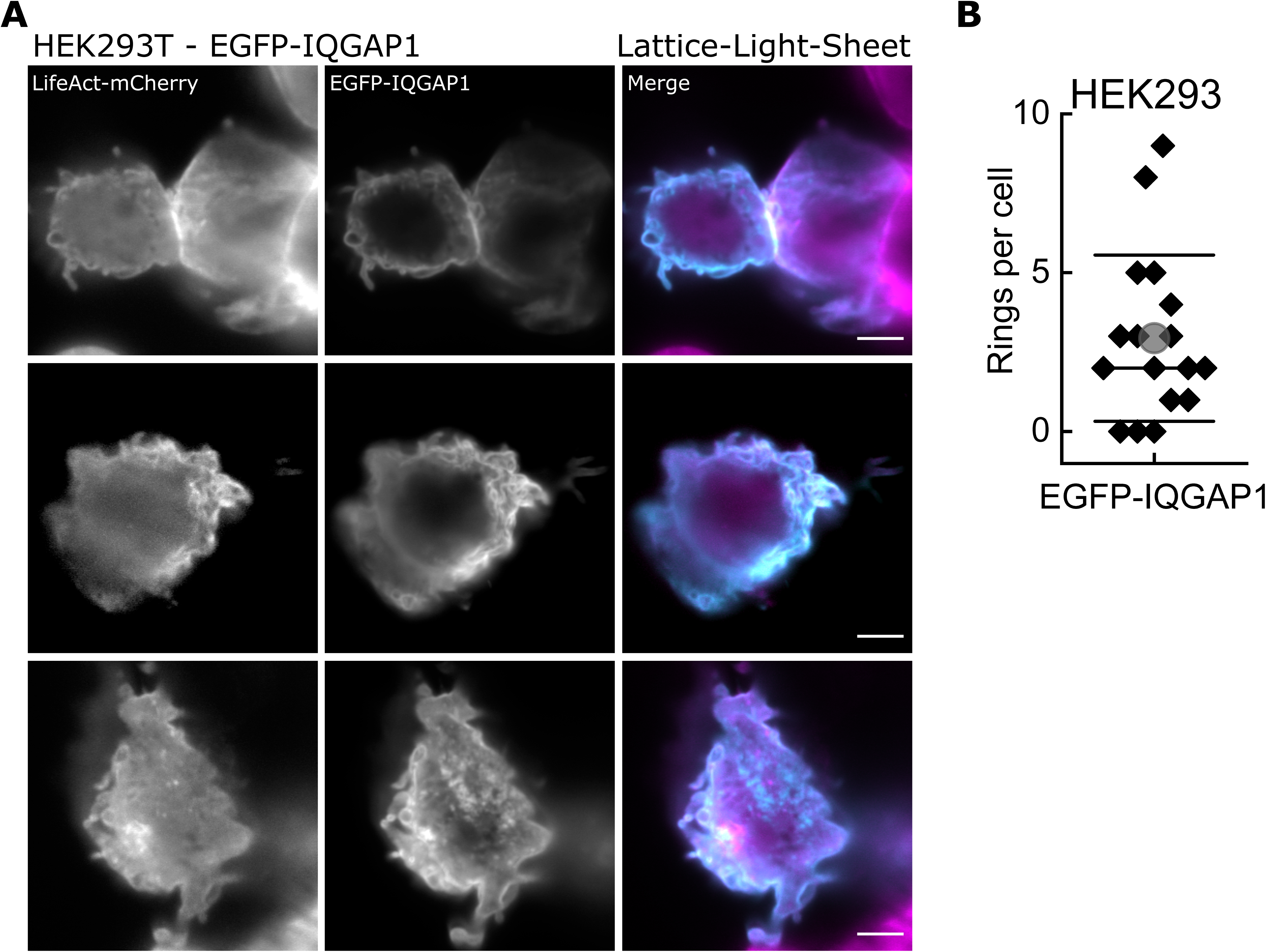
(A) Lattice-light-sheet microscopy images (maximum intensity projection) of HEK293T cells transfected with LifeAct-mCherry (magenta) and EGFP-IQGAP1 (cyan); scale bar: 5 μm. (B) Quantification of actin rings found in HEK293T cell expressing EGFP-IQGAP1.

## Video captions

Video 1: TIRF microscopy image sequence of actin filaments (Alexa488) landing on His_6_-curly decorated SLBs; scale bar: 5 μm.

Video 2: Example image sequence of an actin filament (Alexa488) bound to a His_6_-curly decorated SLB displaying individual bending events after processing the image sequence with a Sobel filter to highlight the shape of the actin filament (the unprocessed images are shown in Figure 2-figure supplement 2); scale bar: 1 μm.

Video 3: Example image sequence of an actin filament (Alexa488, gray) with the plus end labelled by capping protein (SNAP647, yellow) landing on a His_6_-curly decorated SLB; scale bar: 1 μm.

Video 4: Example image sequence of an actin filament (Alexa488, gray) with the plus end labelled by capping protein (SNAP647, cyan) landing on a curly-His_6_ decorated SLB; scale bar: 1 μm.

Video 5: Example image sequences of actin filaments (Alexa488) polymerized by SLB tethered formin in the presence of His_6_-curly bound to the SLB; scale bar 1 μm.

Video 6: Example image sequences of actin filaments (Alexa488) decorated with tropomyosin binding to membrane tethered His_6_-curly; scale bar: 1 μm.

Video 7: Example image sequence showing formation, translation, and contraction of actin filament (Alexa488) rings on membrane tethered His_6_-curly after the addition of muscle myosin II filaments; scale bar: 1 μm.

Video 8: Example image sequences of actin filament (Alexa488) ring contraction on membrane tethered His_6_-curly after the addition of muscle myosin II filaments; scale bar: 1 μm.

Video 9: 3D projection of HEK293T cell expressing EGFP-IQGAP1 (cyan) and Lifeact-mCherry (magenta); images obtained by lattice-light-sheet microscopy; scale bar: 10 μm.

## Notes

### Competing Interest Statement

The authors have declared no competing interest.

### Summary of Updates

implemented changes as suggested by the reviewer comments.

## References

Bieling, P., Li, T.-D., Weichsel, J., McGorty, R., Jreij, P., Huang, B., Fletcher, D. A., & Mullins, R. D. (2016). Force Feedback Controls Motor Activity and Mechanical Properties of Self-Assembling Branched Actin Networks. Cell, 164(1–2), 115–127. https://doi.org/10.1016/j.cell.2015.11.057

Briggs, M. W., & Sacks, D. B. (2003). IQGAP proteins are integral components of cytoskeletal regulation. EMBO Reports, 4(6), 571–574. https://doi.org/10.1038/sj.embor.embor867

De La Cruz, E. M., & Gardel, M. L. (2015). Actin mechanics and fragmentation. Journal of Biological Chemistry, 290(28), 17137–17144. https://doi.org/10.1074/jbc.R115.636472

Eng, K., Naqvi, N. I., Wong, K. C. Y., & Balasubramanian, M. K. (1998). Rng2p, a protein required for cytokinesis in fission yeast, is a component of the actomyosin ring and the spindle pole body. Current Biology, 8(11), 611–621. https://doi.org/10.1016/S0960-9822(98)70248-9

Epp, J. A., & Chant, J. (1997). An IQGAP-related protein controls actin-ring formation and cytokinesis in yeast. Current Biology, 7(12), 921–929. https://doi.org/10.1016/S0960-9822(06)00411-8

Hayakawa, Y., Takaine, M., Imai, T., Yamada, M., Hirose, K., Tokuraku, K., Ngo, K. X., Kodera, N., Numata, O., Nakano, K., & Uyeda, T. Q. P. (2020). Actin binding domain of Rng2 strongly inhibits actin movement on myosin II HMM through structural changes of actin filaments. BioarXiv. https://doi.org/10.1101/2020.04.14.04104

Köster, D. V., Husain, K., Iljazi, E., Bhat, A., Bieling, P., Mullins, R. D., Rao, M., & Mayor, S. (2016). Actomyosin dynamics drive local membrane component organization in an in vitro active composite layer. Proceedings of the National Academy of Sciences, 113(12), E1645–E1654. https://doi.org/10.1073/pnas.1514030113

Kruppa, A. J., Kishi-Itakura, C., Masters, T. A., Rorbach, J. E., Grice, G. L., Kendrick-Jones, J., Nathan, J. A., Minczuk, M., & Buss, F. (2018). Myosin VI-Dependent Actin Cages Encapsulate Parkin-Positive Damaged Mitochondria. Developmental Cell, 44(4), 484–499.e6. https://doi.org/10.1016/j.devcel.2018.01.007

Kučera, O., Janda, D., Siahaan, V., Dijkstra, S. H., Pilátová, E., Zatecka, E., Diez, S., Braun, M., & Lansky, Z. (2020). Anillin propels myosin-independent constriction of actin rings. BioRxiv, 1–27. https://doi.org/10.1101/2020.01.22.915256

Kumari, A., Kesarwani, S., Javoor, M. G., Vinothkumar, K. R., & Sirajuddin, M. (2020). Structural insights into actin filament recognition by commonly used cellular actin markers. The EMBO Journal, 846337. https://doi.org/10.15252/embj.2019104006

Laplante, C., Huang, F., Tebbs, I. R., Bewersdorf, J., & Pollard, T. D. (2016). Molecular organization of cytokinesis nodes and contractile rings by super-resolution fluorescence microscopy of live fission yeast. Proceedings of the National Academy of Sciences, 113(40), E5876–E5885. https://doi.org/10.1073/pnas.1608252113

Laporte, D., Coffman, V. C., Lee, I. J., & Wu, J. Q. (2011). Assembly and architecture of precursor nodes during fission yeast cytokinesis. Journal of Cell Biology, 192(6), 1005–1021. https://doi.org/10.1083/jcb.201008171

Lippincott, J., & Li, R. (1998). Sequential assembly of myosin II, an IQGAP-like protein, and filamentous actin to a ring structure involved in budding yeast cytokinesis. Journal of Cell Biology, 140(2), 355–366. https://doi.org/10.1083/jcb.140.2.355

Litschel, T., Kelley, C. F., Holz, D., Koudehi, M. A., Vogel, S. K., Burbaum, L., Mizuno, N., Vavylonis, D., & Schwille, P. (2020). Reconstitution of contractile actomyosin rings in vesicles. BioRxiv. https://doi.org/10.1101/2020.06.30.180901

Mavrakis, M., Azou-Gros, Y., Tsai, F.-C., Alvarado, J., Bertin, A., Iv, F., Kress, A., Brasselet, S., Koenderink, G. H., & Lecuit, T. (2014). Septins promote F-actin ring formation by crosslinking actin filaments into curved bundles. Nature Cell Biology, 16(4), 322–334. https://doi.org/10.1038/ncb2921

Mccullough, B. R., Blanchoin, L., Martiel, J.-L., De la Cruz, E. M., Cruz, E. M. D. La, & Haven, N. (2008). Cofilin increases the bending flexibility of actin filaments: implications for severing and cell mechanics. Journal of Molecular Biology, 381(3), 550–558. https://doi.org/10.1016/j.jmb.2008.05.055

Mishra, M., Kashiwazaki, J., Takagi, T., Srinivasan, R., Huang, Y., Balasubramanian, M. K., & Mabuchi, I. (2013). In vitro contraction of cytokinetic ring depends on myosin II but not on actin dynamics. Nature Cell Biology, 15(7), 853–859. https://doi.org/10.1038/ncb2781

Miyazaki, M., Chiba, M., Eguchi, H., Ohki, T., & Ishiwata, S. (2015). Cell-sized spherical confinement induces the spontaneous formation of contractile actomyosin rings in vitro. Nature Cell Biology, 17(4), 480–489. https://doi.org/10.1038/ncb3142

Mosby, L. S., Hundt, N., Young, G., Fineberg, A., Polin, M., Mayor, S., Kukura, P., & Köster, D. V. (2020). Myosin II Filament Dynamics in Actin Networks Revealed with Interferometric Scattering Microscopy. Biophysical Journal, 118(8), 1946–1957. https://doi.org/10.1016/j.bpj.2020.02.025

Murrell, M. P., & Gardel, M. L. (2012). F-actin buckling coordinates contractility and severing in a biomimetic actomyosin cortex. Proceedings of the National Academy of Sciences of the United States of America, 109(51), 20820–20825. https://doi.org/10.1073/pnas.1214753109

Nye, J. A., & Groves, J. T. (2008). Kinetic control of histidine-tagged protein surface density on supported lipid bilayers. Langmuir_: The ACS Journal of Surfaces and Colloids, 24(8), 4145–4149. https://doi.org/10.1021/la703788h

Padmanabhan, A., Bakka, K., Sevugan, M., Naqvi, N. I., D’Souza, V., Tang, X., Mishra, M., & Balasubramanian, M. K. (2011). IQGAP-related Rng2p organizes cortical nodes and ensures position of cell division in fission yeast. Current Biology, 21(6), 467–472. https://doi.org/10.1016/j.cub.2011.01.059

Palani, S., Köster, D. V., Hatano, T., Kamnev, A., Kanamaru, T., Brooker, H. R., Hernandez-Fernaud, J. R., Jones, A. M. E., Millar, J. B. A., Mulvihill, D. P., & Balasubramanian, M. K. (2019). Phosphoregulation of tropomyosin is crucial for actin cable turnover and division site placement. The Journal of Cell Biology, 218(11), 3548–3559. https://doi.org/10.1083/jcb.201809089

Pardee, J. D., & Spudich, J. A. (1982). Purification of muscle actin. Methods in Cell Biology, 24, 271–289. http://www.ncbi.nlm.nih.gov/pubmed/7098993

Schmid, B., Tripal, P., Fraaß, T., Kersten, C., Ruder, B., Grüneboom, A., Huisken, J., & Palmisano, R. (2019). 3Dscript: animating 3D/4D microscopy data using a natural-language-based syntax. Nature Methods, 16(4), 278–280. https://doi.org/10.1038/s41592-019-0359-1

Singh, S. M., Bandi, S., & Mallela, K. M. G. (2017). The N-Terminal Flanking Region Modulates the Actin Binding Affinity of the Utrophin Tandem Calponin-Homology Domain. Biochemistry, 56(20), 2627–2636. https://doi.org/10.1021/acs.biochem.6b01117

Skau, C. T., & Kovar, D. R. (2010). Fimbrin and tropomyosin competition regulates endocytosis and cytokinesis kinetics in fission yeast. Current Biology, 20(16), 1415–1422. https://doi.org/10.1016/j.cub.2010.06.020

Skoumpla, K., Coulton, A. T., Lehman, W., Geeves, M. A., & Mulvihill, D. P. (2007). Acetylation regulates tropomyosin function in the fission yeast Schizosaccharomyces pombe. Journal of Cell Science, 120(9), 1635–1645. https://doi.org/10.1242/jcs.001115

Taylor, K. A., Taylor, D. W., & Schachat, F. (2000). Isoforms of α-actinin from cardiac, smooth, and skeletal muscle form polar arrays of actin filaments. Journal of Cell Biology, 149(3), 635–645. https://doi.org/10.1083/jcb.149.3.635

Tebbs, I. R., & Pollard, T. D. (2013). Separate roles of IQGAP Rng2p in forming and constricting the Schizosaccharomyces pombe cytokinetic contractile ring. Molecular Biology of the Cell, 24(12), 1904–1917. https://doi.org/10.1091/mbc.E12-10-0775

Vassilopoulos, S., Gibaud, S., Jimenez, A., Caillol, G., & Leterrier, C. (2019). Ultrastructure of the axonal periodic scaffold reveals a braid-like organization of actin rings. Nature Communications, 10(1), 636217. https://doi.org/10.1038/s41467-019-13835-6

Wachsstock, D. H., Schwartz, W. H., & Pollard, T. D. (1993). Affinity of alpha-actinin for actin determines the structure and mechanical properties of actin filament gels. Biophysical Journal, 65(1), 205–214. https://doi.org/10.1016/S0006-3495(93)81059-2

Wang, C.-H., Balasubramanian, M. K., & Dokland, T. (2004). Structure, crystal packing and molecular dynamics of the calponin-homology domain of Schizosaccharomyces pombe Rng2. Acta Crystallographica Section D Biological Crystallography, 60(8), 1396–1403. https://doi.org/10.1107/S0907444904012983

Way, M., Sanders, M., Garcia, C., Sakai, J., & Matsudaira, P. (1995). Sequence and domain organization of scruin, an actin-cross-linking protein in the acrosomal process of Limulus sperm. Journal of Cell Biology, 128(1–2), 51–60.

Xu, K., Zhong, G., & Zhuang, X. (2013). Actin, spectrin, and associated proteins form a periodic cytoskeletal structure in axons. Science (New York, N. Y.), 339(6118), 452–456. https://doi.org/10.1126/science.1232251

